# A multi-scale digital twin for adiposity-driven insulin resistance in humans: diet and drug effects

**DOI:** 10.1101/2023.04.20.537480

**Authors:** Tilda Herrgårdh, Christian Simonsson, Mattias Ekstedt, Peter Lundberg, Karin G. Stenkula, Elin Nyman, Peter Gennemark, Gunnar Cedersund

## Abstract

**Aims:** The increased prevalence of insulin resistance is one of the major health risks in society today. Insulin resistance involves both short-term dynamics, such as altered meal responses, and long-term dynamics, such as development of type 2 diabetes. Insulin resistance also occurs on different physiological levels, ranging from disease phenotypes to organ-organ communication and intracellular signaling. To better understand the progression of insulin resistance, an analysis method is needed that can combine different timescales and physiological levels. One such method is digital twins, consisting of combined mechanistic multi-scale and multi-level mathematical models. We have previously developed a multi-level model for short-term glucose homeostasis and intracellular insulin signaling, and there exists long-term weight regulation models. However, no one has combined these kinds of models into an interconnected, multi-level and multi-timescale digital twin model. Herein, we present a first such multi-scale digital twin for the progression of insulin resistance in humans.

**Methods:** The model is based on ordinary differential equations representing biochemical and physiological processes, in which unknown parameters were fitted to data using a MATLAB toolbox.

**Results:** The connected twin correctly predicts independent data from a weight increase study, both for weight-changes, for fasting plasma insulin and glucose levels, as well as for intracellular insulin signaling. Similarly, the model can predict independent weight-change data in a weight loss study, involving diet and the weight loss drug topiramate. These independent validation tests are confirmed by a chi-square test (𝑉(𝜃) = 4.8 < 21 = 𝜒^2^_𝑐𝑢𝑚,𝑖𝑛𝑣_ (12,0.05)). In both these cases, the model can also predict non-measured variables, such as activity of intracellular intermediaries, glucose tolerance responses, and organ fluxes.

**Conclusions:** We present a first multi-level and multi-timescale model, describing dynamics on the whole-body, organ and cellular levels, ranging from minutes to years. This model constitutes the basis for a new digital twin technology, which in the future could potentially be used to aid medical pedagogics and increase motivation and compliance and thus aid in prevention and treatment of insulin resistance.

## Introduction

Insulin resistance is becoming more common, partly due to a general weight increase in the population, and it is one of today’s major health problems. Insulin resistance is both a part of, and a precursor of, type 2 diabetes. The progression towards these harmful conditions is complex: they usually develop over many years, involving both short and long-term changes with dynamics ranging from minutes to years. Furthermore, the changes happen on different biological levels: inside cells, within and between organs, and on the whole-body level. A widely spread hypothesis for the cause of type 2 diabetes is adiposity-driven insulin resistance: an impaired or saturated lipid storage capacity in adipose tissue cause ectopic accumulation of lipids on other organs and tissues, for example in muscle tissue and liver. This lipid accumulation is associated with decreased insulin sensitivity in insulin’s target tissues. This decreased insulin sensitivity is initially compensated for by increased insulin secretion, but over time leads to pancreatic failure. The resulting reduction in insulin secretion leads to the onset of type 2 diabetes (1). It is therefore clear that the progression of insulin resistance is a complex, multi-level, multi-timescale process.

Some of the processes and risk factors for insulin resistance, such as your genetic risk and age, are not controllable. Still, both insulin resistance and type 2 diabetes are preventable, manageable, and possibly even treatable. Regarding prevention, maintaining a low weight is viewed as one of the most important strategies. Regarding treatment, it has recently been shown that weight reduction is sometimes able to reverse type 2 diabetes (2). For some individuals, weight reduction might be more difficult for a variety of reasons, and in these cases, a weight reducing drug might be valuable. The choice of drug for prevention, management, and/or treatment is complex due to the inherent heterogeneity in type 2 diabetes in different individuals, and due to the varying effects of different drugs and diet/exercise regimes. This complexity in treatment choices, as well as the multi-scale complexity in disease mechanisms, points to a need for a more comprehensive understanding of insulin resistance, both on a general and an individual level. One method for achieving, testing, and visualizing such a comprehensive understanding is to represent this understanding using mathematical models and digital twins.

Digital twins and mechanistic modelling have been used extensively to study different individual aspects of the insulin resistance and type 2 diabetes, on both whole-body, organ or tissue, and cellular level. For whole-body weight regulation, there exists models that describe body composition as a response to energy intake, such as the one developed by Hall et al (3). For the organ and tissue level, meal response models such as that developed by Dalla Man et al. are relevant, and have even been approved by the US Food and Drug Administration (FDA) for certain applications (4,5). On the cellular level, there exists models that describe e.g. pancreas, liver, and adipocytes (6–8). There also exist some models that combine these different levels in comprehensive model that can explain both short- and long-term dynamics. Such multi-level models include the longitudinal model developed by Ha et al. (9), that describes two different progressions towards type 2 diabetes. Another model, developed by Uluseker et al. (10), combines the Dalla Man model with an adipocyte model. We have also developed such a multi-level model, combining an adipocyte model for intracellular insulin signaling with the Dalla Man model for organ-organ communication in glucose homeostasis (4). However, to the best of our knowledge there exists no multi-level and multi-timescale model that can describe data for all three levels, and that can describe the progression into diabetes in a mechanistic manner.

Herein, we present a first multi-level, multi-timescale, and mechanistic mathematical model that also can describe the progression to diabetes in a semi-mechanistic manner (Fig. 1B). The model uses a new adiposity-driven insulin resistance model to connect the three different physiological levels and timescales: long-term whole-body weight composition over months and years; short-term meal-responses of glucose and insulin over a few hours, and fast dynamics of adipocyte intracellular insulin signaling over seconds and minutes. We develop and test the model using data from two scenarios: i) the progression towards insulin resistance due to weight gain, where the model correctly predicts data for fasting glucose and insulin levels, as well as intracellular insulin signaling in adipocytes (Fig. 1C), and ii) a weight loss scenario, where the model can describe data for both weight-loss due to decreased energy intake alone, and due to additional drug usage (Fig. 1D). We also use the model to make predictions regarding relevant biomarkers for type 2 diabetes not measured in the above studies, showcasing how the model can be used to unravel more processes than those directly measured. Future iterations of the model could potentially be used to evaluate different health scenarios, e.g., different diets and medications, and as such aid in the prevention and treatment of insulin resistance.

**Figure 1.**
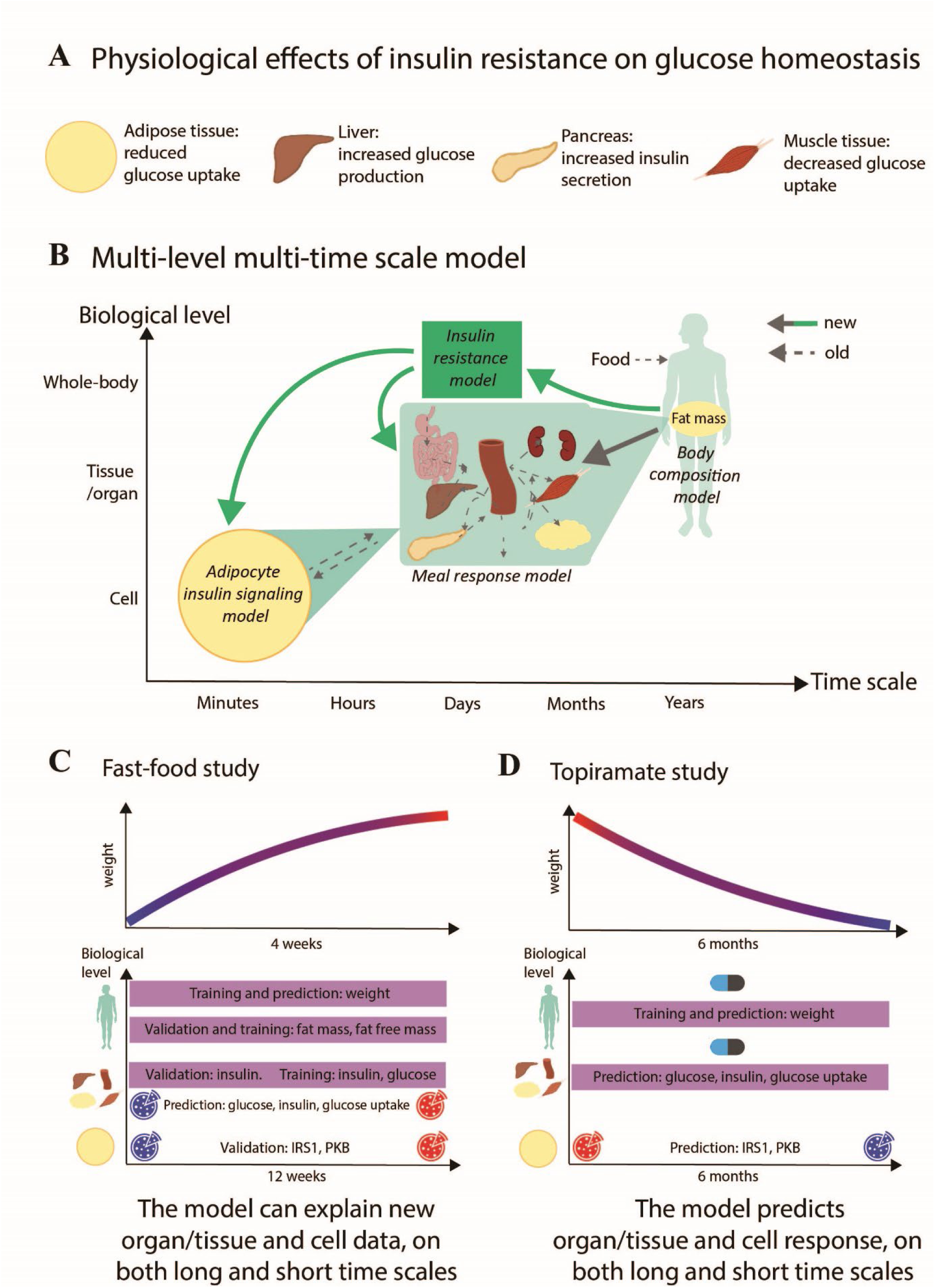
A) The different physiological effects of insulin resistance on glucose homeostasis. B) Schematic overview of the multi-level and multi-scale model structure, connecting multiple body levels and timescales. The new reactions (solid lines) include a connection from the Body composition model on the whole-body level to the Meal response model on the organ/tissue level, and the intracellular level, as well as arrows to and from the new insulin resistance model. Reactions in previously published models are shown as dashed lines. C) Schematic overview of the analyses made herein, involving two different studies: a weight gain study – the Fast-food study – that was conducted during 12 weeks, and a weight decrease study on the drug topiramate – the Topiramate study. Fast-food study: on the whole-body level, the model was trained on weight data as well as fat mass and fat-free mass data, and validated on fat mass and fat free mass, as shown in detail in Fig. 3BC and Fig. 3C respectively. A prediction of further weight increase was also made, shown in Fig. 4A. On the organ/tissue level, the model was validated on fasting insulin data, shown in Fig. 3D, and predictions were made of meal response insulin, glucose, and glucose uptake in fat and muscle tissue before and after the diet, as shown in Fig. 4B. D) Topiramate study: on the whole-body level, the model was trained and validated on weight data for placebo and 3 different dosages of Topiramate. The model was then used to predict two other scenarios not explored in the study – an increase in energy intake with and without medication – as well as meal responses before and after these scenarios, on both organ/tissue and cell level.

## Method

### Model equations

The models are built up by standard form ordinary differential equations (ODEs). All of equations are given in the supplementary material, both as equations and as simulation files. Below we only describe the equations that were added to the multi-level model in this article, specifically those of the *insulin resistance model*, the *weight-meal response interconnection*, the *phenomenological energy intake*, and the *drug response model for topiramate*.

### Insulin resistance on organ/tissue level

The insulin resistance part of the model is inspired by the similar insulin resistance equations implemented for mice in (11). The equations used herein are:

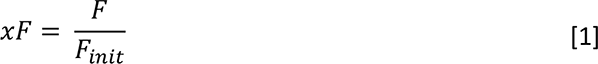

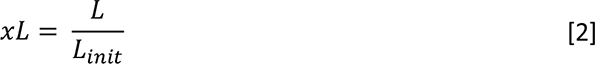

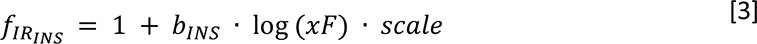

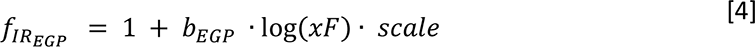

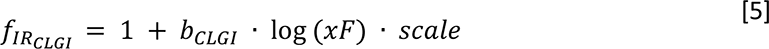

where 𝑥𝐹 is the relative change in fat mass from initial fat mass 𝐹_𝑖𝑛𝑖𝑡_, 𝐹 is the current fat mass, 𝑥𝐿 is the relative change in lean tissue mass from initial lean tissue mass 𝐿_𝑖𝑛𝑖𝑡_, 𝐿 is the current lean tissue mass, 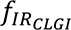 is the insulin resistance effect on hepatic and muscle glucose uptake, 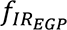 is the insulin resistance effect on endogenous glucose production, 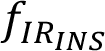 is the insulin resistance effect on insulin secretion, 𝑠𝑐𝑎𝑙𝑒 scales all the insulin resistance effects from mice to humans, and 𝑏_𝐶𝐿𝐺𝐼_, 𝑏_𝐸𝐺𝑃_, 𝑏_𝐼𝑁𝑆_ are parameters.

The effect of the insulin resistance on insulin secretion is described by

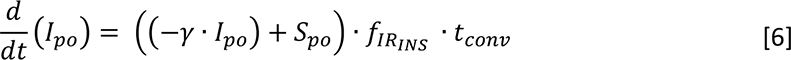

where 𝐼_𝑝𝑜_ is the amount of insulin in the portal vein, 𝛾 is the transfer rate constant between portal vein and liver, 𝑆_𝑝𝑜_ is the insulin secretion into the portal vein, 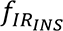 is the insulin resistance effect on insulin secretion, and 𝑡_𝑐𝑜𝑛𝑣_ is a parameter for time conversion between the body composition model, defined in the time units days, and the other models, defined in minutes. Note that this is different from the previously reported mouse model, where the insulin resistance effect is directly on 𝑆_𝑝𝑜_. One of the reasons for this difference is that insulin in the portal vein, 𝐼_𝑝𝑜_, is not explicitly modelled in the mouse model.

The effect of insulin resistance on endogenous glucose production, 𝐸𝐺𝑃, is described by

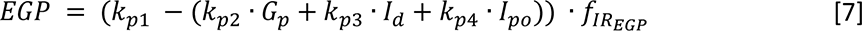

where 𝑘_𝑝1_ is the extrapolated 𝐸𝐺𝑃 at zero glucose and insulin, 𝑘_𝑝2_ is liver glucose effectiveness, 𝑘_𝑝3_ governs the amplitude of insulin action on the liver, 𝐼_𝑑_ is a delayed insulin signal, 𝑘_𝑝4_ governs the amplitude of portal insulin action on the liver.

The effect of insulin resistance on glucose utilization in the liver, 𝑈_𝑖𝑑𝑙_, and muscle tissue, 𝑈_𝑖𝑑𝑚_, is affected by insulin resistance as follows:

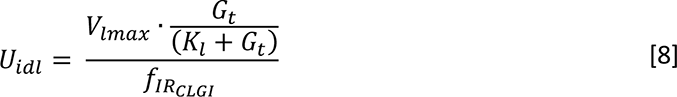

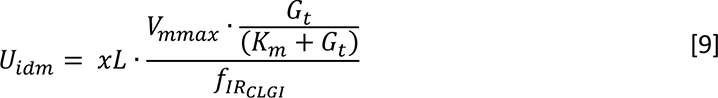

where 𝑉_𝑙𝑚𝑎𝑥_ is the maximum rate of glucose utilization in the liver, 𝐺_𝑡_ is the glucose in tissue, 𝐾_𝑙_ is a Michaelis-Menten parameter, 𝑉_𝑚𝑚𝑎𝑥_ is the maximum rate of glucose utilization in muscle, and 𝐾_𝑚_ is a Michaelis-Menten parameter. In our model, insulin resistance does not directly influence the glucose utilization in fat tissue, 𝑈_𝑖𝑑_**_𝑓_** since it has been observed in diabetics that glucose uptake is significantly changed in muscle and liver but not in fat tissue. (12,13).

### Insulin resistance on cell level

The insulin resistance on the cell level is implemented as a gradual transition between the different parameter sets for non-diabetics and diabetics from the previous version of the model. The effect of diabetes was, as in the previous model, implemented on three different places in the model: 𝐼𝑅, 𝐺𝐿𝑈𝑇4, and 𝑑𝑖𝑎𝑏𝑒𝑡𝑒𝑠. The diabetes effect on 𝐼𝑅 decreases the total amount of 𝐼𝑅, and with less insulin receptors, less insulin can bind to the cell, i.e. the cell is less sensitive to insulin. The diabetes effect on 𝐺𝐿𝑈𝑇4 decreases the amount of 𝐺𝐿𝑈𝑇4, which means that less 𝐺𝐿𝑈𝑇4, can be taken up by the cell. The parameter named 𝑑𝑖𝑎𝑏𝑒𝑡𝑒𝑠 reduces the positive feedback from 𝑚𝑇𝑂𝑅𝐶1 to 𝐼𝑅𝑆1 (Fig. 3E). All these diabetes effects results in an increase in insulin sensitivity and a decrease in glucose uptake in the model. The gradual transition of these diabetes effects was, as with previous insulin resistance equations, dependent on the change in fat mass as follows:

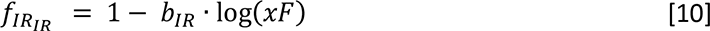

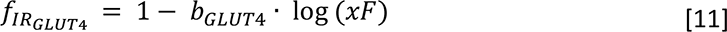

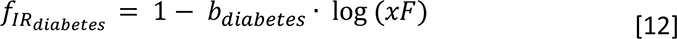

where 𝑏_𝐼𝑅_, 𝑏_𝐺𝐿𝑈𝑇4_, and 𝑏_𝑑𝑖𝑎𝑏𝑒𝑡𝑒𝑠_ are parameters.

As mentioned, the diabetes effect was static in the previous model – the model could either be diabetic, non-diabetic, but could not transition from one to the other. A transition between non-diabetic and diabetic version of the model was not possible since the total amount of 𝐼𝑅 and 𝐺𝐿𝑈𝑇4 could not change. To make the gradual transition to diabetes possible, equations that could change the total amount of 𝐼𝑅 and 𝐺𝐿𝑈𝑇4 was therefore added. Specifically, degradation and protein expression of 𝐼𝑅 and 𝐺𝐿𝑈𝑇4 was added (Fig. 4C). The protein expressions of 𝐼𝑅 and 𝐺𝐿𝑈𝑇4 are then influenced by the insulin resistance functions 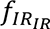 and 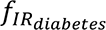 to achieve the gradual decrease of 𝐼𝑅 and 𝐺𝐿𝑈𝑇4 that is part of the gradual transition to diabetes (Eq. 15 and 19). For 𝐼𝑅, the following equations were changed:

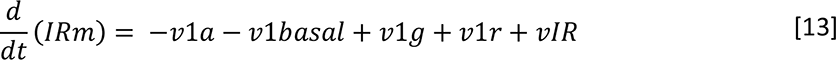

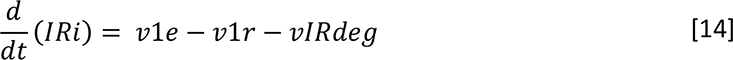

where 𝐼𝑅𝑚 is the insulin receptors (𝐼𝑅) found in the cell membrane, 𝑣1𝑎, 𝑣1𝑏𝑎𝑠𝑎𝑙, 𝑣1𝑔, and 𝑣1𝑟 are the unchanged reaction rates describing the transition of 𝐼𝑅𝑚 to and from other 𝐼𝑅-forms (see the supplementary material and (7)), and 𝑣𝐼𝑅 is the new reaction rate describing the protein expression of 𝐼𝑅𝑚, 𝐼𝑅𝑖𝑑 is the internalized form of 𝐼𝑅, 𝑣1𝑒 and 𝑣1𝑟 are the unchanged reaction rates describing the transition of 𝐼𝑅𝑖 to and from other 𝐼𝑅-forms (see the supplementary material and (7)), 𝑣𝐼𝑅 is the new reaction rate describing the protein expression of 𝐼𝑅𝑚, and 𝑣𝐼𝑅𝑑𝑒𝑔 is the new reaction rate describing the degradation of 𝐼𝑅𝑖. The reaction rates 𝑣𝐼𝑅𝑑𝑒𝑔 and 𝑣𝐼𝑅 are defined as:

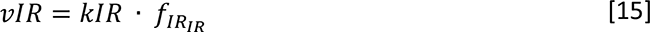

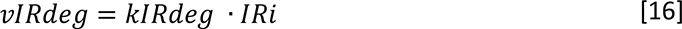

where 𝑘𝐼𝑅 and 𝑘𝐼𝑅𝑑𝑒𝑔 are parameters. For 𝐺𝐿𝑈𝑇4, the following equations where changed:

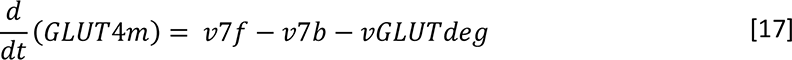

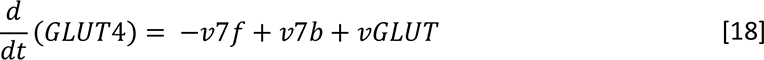

where 𝐺𝐿𝑈𝑇4𝑚 and 𝐺𝐿𝑈𝑇4 are the two forms of 𝐺𝐿𝑈𝑇4, the first associated with the cellular membrane and the other inside the cell cytosol, 𝑣7𝑓 and 𝑣7𝑏 are the unchanged reaction rates describing the transition between 𝐺𝐿𝑈𝑇4𝑚 and 𝐺𝐿𝑈𝑇4, 𝑣𝐺𝐿𝑈𝑇𝑑𝑒𝑔 is the new reaction describing the degradation of 𝐺𝐿𝑈𝑇4𝑚, and 𝑣𝐺𝐿𝑈𝑇 is the new reaction rate describing the protein expression of 𝐺𝐿𝑈𝑇4.

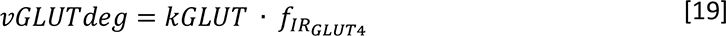

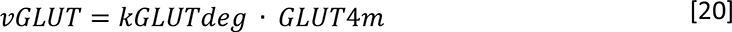

where 𝑘𝐺𝐿𝑈𝑇 and 𝑘𝐺𝐿𝑈𝑇𝑑𝑒𝑔 are parameters. The membrane form, 𝐺𝐿𝑈𝑇4𝑚, then effects the inflow of glucose to the cell, which is upscaled to 𝑈_𝑖𝑑_**_𝑓_** as described in (14).

The now gradual adiposity driven effect of insulin resistance on the positive feedback from 𝑚𝑇𝑂𝑅𝐶1 to 𝐼𝑅𝑆1, 𝑣2𝑐, was applied in the same way as the parameter 𝑑𝑖𝑎𝑏𝑒𝑡𝑒𝑠 was in the previous model:

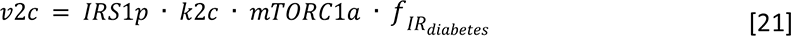

where 𝐼𝑅𝑆1𝑝 is the amount of phosphorylated form of 𝐼𝑅𝑆1, 𝑘2𝑐 is a parameter, 𝑚𝑇𝑂𝑅𝐶1𝑎 is the amount of 𝑚𝑇𝑂𝑅𝐶1𝑎.

### Weight-meal response interconnection

As shown in Equation 9, the change in lean tissue mass, 𝑥𝐿, has a direct effect on the glucose utilization in muscle tissue. This effect is a part of the connection between the whole-body weight model and the meal response model. The glucose utilization in fat tissue, 𝑈_𝑖𝑑_**_𝑓_**, is also affected by the weight model, specifically by the change in fat mass:

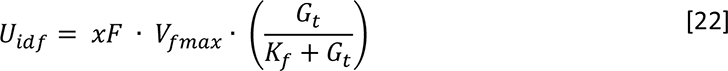

where, similarly to the utilization in the other tissues, 𝑉**_𝑓_**_𝑚𝑎𝑥_ is the maximum rate of glucose utilization in muscle, and 𝐾**_𝑓_** is a Michaelis-Menten parameter.

Equations 9 and 22 also show the connection between the whole-body and the organ/tissue level: the glucose uptake in muscle and fat tissue changes with the change in lean and fat mass respectively. Furthermore, the glucose rate of appearance, 𝑅𝑎, changes with the total body weight (𝐵):

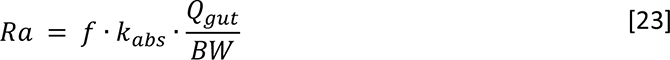

where 𝑓 is the fraction of intestinal glucose absorption which appears in plasma, 𝑘_𝑎𝑏𝑠_ is the absorption rate, 𝑄_𝑔𝑢𝑡_ is the glucose content in the gut, and 𝐵𝑊 is the body weight. In the earlier model, 𝐵𝑊 was a constant, while here it is a variable in the whole-body level as described in (3).

To merge the different models, a parameter 𝑡_𝑐𝑜𝑛𝑣_ = 24 · 60 was introduced to the models corresponding to time expressed in minutes, i.e., the organ/tissue level model and the cell model, to change the unit for time into days.

### Phenomenological energy intake

We added an equation for accounting for differences in energy intake throughout the study period:

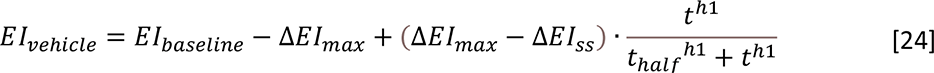

where 𝐸𝐼_𝑣𝑒ℎ𝑖𝑐𝑙𝑒_ (𝑡) is the energy intake over time, 𝐸𝐼_𝑏𝑎𝑠𝑒𝑙𝑖𝑛𝑒_ is the energy intake at baseline, Δ𝐸𝐼_𝑚𝑎𝑥_ is the maximum change in energy intake, here fixed at the change in energy intake that the participants were asked to follow, Δ𝐸𝐼_𝑠𝑠_ is the change in energy intake at steady state, 𝑡 is the time, ℎ1 is the hill coefficient, and 𝑡_ℎ𝑎𝑙_**_𝑓_** is the timepoint where half of 𝐸𝐼_𝑣𝑒ℎ𝑖𝑐𝑙𝑒_(𝑡) has been reached.

### Drug response model for topiramate

The energy intake was also altered with respect to the drug topiramate according to

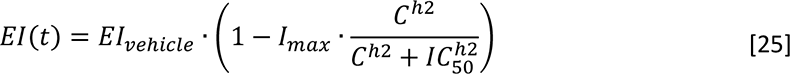

where 𝐸𝐼(𝑡) is the energy intake that influenced by topiramate, ℎ2, 𝐼_𝑚𝑎𝑥_, and 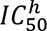 are parameters, and 𝐶 is the concentration of topiramate in plasma. To get 𝐶, we adopted the standard two-compartment pharmacokinetic model with first-order absorption from (15)

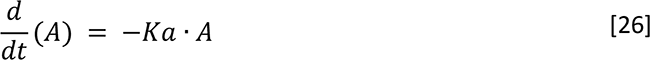

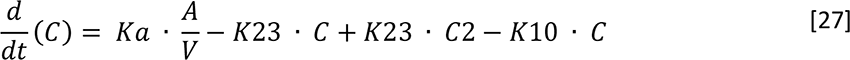

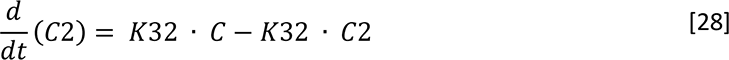

where 𝐴 is the absorption compartment, into which the daily dosages of topiramate are administered, 𝐾𝑎, 𝑉, 𝐾23, 𝐾10, and 𝐾32 are parameters, and 𝐶2 is the topiramate concentration in tissue.

### Parameter Estimation

Almost all of the 146 parameters in this multi-scale model were fixed at their values obtained from previous studies. The parameters estimated in this article are one scaling parameter of the insulin resistance model, the scaling parameter of the diabetes effects in the cell-level model, the new parameters in the cell-level model, those parameters corresponding to the phenomenological energy intake equation, and finally the parameters of the meal response model. The different parameters are estimated using different data and in different ways.

Most parameters were optimized using an optimization algorithm. Specifically, the parameters were estimated by minimizing the difference between model simulations, denoted 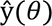, and experimental data, denoted 𝑦. The cost function used is the conventional weight least square, i.e.,

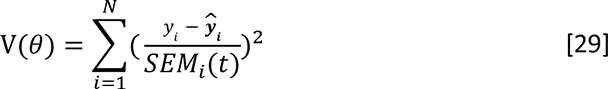

where the subscript 𝑖 denotes the data point, where 𝑁 denotes the number of data points, and where 𝑆𝐸𝑀 denotes the standard error of the mean for the data uncertainty (15). In practice, this parameter estimation was accomplished using the enhanced scatter search (eSS) algorithm from the MEIGO toolbox (16). The optimization was restarted multiple times, run in parallel at the local node of the Swedish national supercomputing center (NSC). The parameter estimation was allowed to freely find the best possible combinations of parameter values within boundaries.

We use a 𝜒^2^-test to evaluate the agreement between model simulations and data. To be more specific, we use the inverse of the cumulative 𝜒^2^-distribution function for setting a threshold, 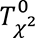, and then compare the cost function 𝑉(𝜃) with this threshold:

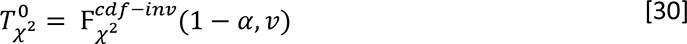

where 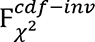 us the inverse density function, 𝛼 is the significance level, and 𝑣 is the degrees of freedom, which was the same as the number of data points in the training data sets. The model is then rejected if the model cost is larger than 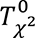 .

The parameters that were not estimated using an algorithm were estimated manually due to simplicity, but the fit to data was assessed in the same way as for the optimization algorithm, i.e. with a 𝜒^2^-test (Eq. 30). Note that apart from these explicitly mentioned parameters, all other parameters were optimized using an optimization algorithm (Eq. 29).

The scaling parameter of the insulin resistance model, which accounts for the scale difference in fat tissue between mice and humans, was estimated by hand. The data used for this manual fitting was the fasting insulin data from the Fast-food study (17,18) (Fig. 3D).

The scaling of the three diabetes effects - 𝐼𝑅, 𝐺𝐿𝑈𝑇4, and 𝑑𝑖𝑎𝑏𝑒𝑡𝑒𝑠 - were adjusted by hand to fit to the level of diabetes seen in the cellular data from the Fast-food study (Fig. 3G). The three diabetes effects have their own range of diabetic to non-diabetic values (Fig. 3EF) – 55-100 for 𝐼𝑅, 50-100 for 𝐺𝐿𝑈𝑇4, and 15.5-100 for 𝑑𝑖𝑎𝑏𝑒𝑡𝑒𝑠. These ranges of diabetes effects where then scaled using one scaling parameter, scaling them towards a percentage of diabetes that corresponded to an acceptable fit to the cellular data after the fast-food diet.

The parameters added to the cell model to enable a gradual change due to insulin resistance, 𝑘𝐺𝐿𝑈𝑇, 𝑘𝐺𝐿𝑈𝑇𝑑𝑒𝑔, 𝑘𝐼𝑅 and 𝑘𝐼𝑅𝑑𝑒𝑔 (Eq. 13-20), where also adjusted manually. These parameters were adjusted so that the initial values of total 𝐼𝑅 and total 𝐺𝐿𝑈𝑇4 had a steady state at 100%.

The last parameters to be adjusted manually where the parameters of the insulin resistance equations (Eq. 10-12), 𝑏_𝐼𝑅_, 𝑏_𝐺𝐿𝑈𝑇4_, and 𝑏_𝑑𝑖𝑎𝑏𝑒𝑡𝑒𝑠_. These parameters were adjusted so that the initial values of total 𝐼𝑅 and total 𝐺𝐿𝑈𝑇4 reached the scaled values from the estimation to cellular data within the time span of the Fast-food study (Fig. 4D).

Two sets of parameters were adjusted using an optimization algorithm: the energy intake parameters and the meal response parameters. The parameters relating to the energy-intake equation were estimated using data from the Topiramate study. This estimation data consists of body-weight time-course data, which is denoted 𝐵W. The meal-response parameters were estimated using the baseline values of fasting plasma insulin and glucose from the Fast-food study, and were only changed when used in the training and predictions relating to the Fast-food study (i.e., the training and prediction related to the Topiramate study used the parameters from the original article (4)). These parameters were kept within tight bounds (a factor of 1) of the parameter values from the original model (4).

For detailed description of all parameters, see the supplementary material. All other parameters were fixed and set to values used in Nyman et al. (2011), and these values are listed in the supplementary material. We exploited the modular structure of the model by fitting the weight model on its own. In the final simulation with the multi-level model, all aspects of the model are simulated at the same time.

### Model simulation

We exploited the unidirectional structure of the multi-level model to only simulate those parts of the model that are needed. In other words, the whole-body part of the model is not impacted by other parts of the model and could therefore be simulated on its own, for instance when estimating the parameters in that part of the model to only the weight data. In contrast, the entire multi-level model was simulated for the tissue- and cell-levels.

The initial values used in the simulations can be found in the supplementary material.

### Uncertainty estimation

The uncertainty of both the parameters and the model simulations for estimation, validation, and predictions were gathered as proposed in (19) and as implemented in (20). In short, the desired property (i.e., the fasting plasma glucose and insulin levels in the Fast-food study (Fig. 3) and the weight data in the Topiramate study (Fig. 4)) were either maximized or minimized, while requiring the cost to be below the 𝜒^2^-threshold. See (20) for more details on how the uncertainty estimation was done.

### Model and data availability

We used MATLAB R2020b (MathWorks, Natick, MA) and the IQM toolbox (IntiQuan GmbH, Basel, Switzerland) for the entire modelling work performed (21).

No new experimental data were collected in this study. We therefore refer to the methods sections in the original articles (17,18,22–24) for the corresponding details experimental methods.

## Results

### Mechanistic, multi-level, and multi-timescale model

The multi-level model (Fig. 2) is comprised of three interconnected models, previously published on their own, plus a new insulin resistance model adopted from rodents (11). Firstly, the whole-body model describes changes on body-composition (3), which produces input to the new sub-model for the progression of insulin resistance. Secondly, the tissue-level model describes the meal response of plasma glucose, organ-specific glucose uptake, and insulin regulation (4). Thirdly, the cellular level describes intracellular insulin signaling in adipocytes (6). The whole-body model has previously been trained and validated on weight-change data (3), and the interconnected tissue-level and cell-level model was previously trained and validated on meal-response data and intracellular insulin-signaling data from human adipocytes (6). The insulin resistance model (Fig. 2A, green box) is, as in (11), driven by adiposity, specifically the relative increase in fat mass from baseline 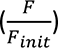. The insulin resistance affects the tissue-level model in three ways: 1) it decreases glucose utilization in muscle and liver tissue (𝑈_𝑖𝑑𝑚_, 𝑈_𝑖𝑑_**_𝑓_**) (Eq. 5), 2) it increases endogenous glucose production (𝐸𝐺𝑃) (Eq. 4), and 3) it increases insulin secretion (𝐼_𝑝𝑜_) (Eq 3). The insulin resistance model also influences the cell-level model in three ways: by decreasing the protein expression of 𝐼𝑅 and 𝐺𝐿𝑈𝑇4, and reducing a positive feedback from 𝑚𝑇𝑂𝑅𝐶1 to 𝐼𝑅𝑆1. The connection between the whole-body model and the tissue-level model is top-down, and comprises three parts: 1) muscle uptake (𝑈_𝑖𝑑𝑚_) is dependent on muscle mass, 2) adipose tissue uptake (𝑈_𝑖𝑑_**_𝑓_**) is dependent on fat free mass, and 3) rate of appearance of glucose (𝑅𝑎) is dependent on total body weight (𝐵W). All model equations and parameter values can be found in the Supplement.

**Figure 2.**
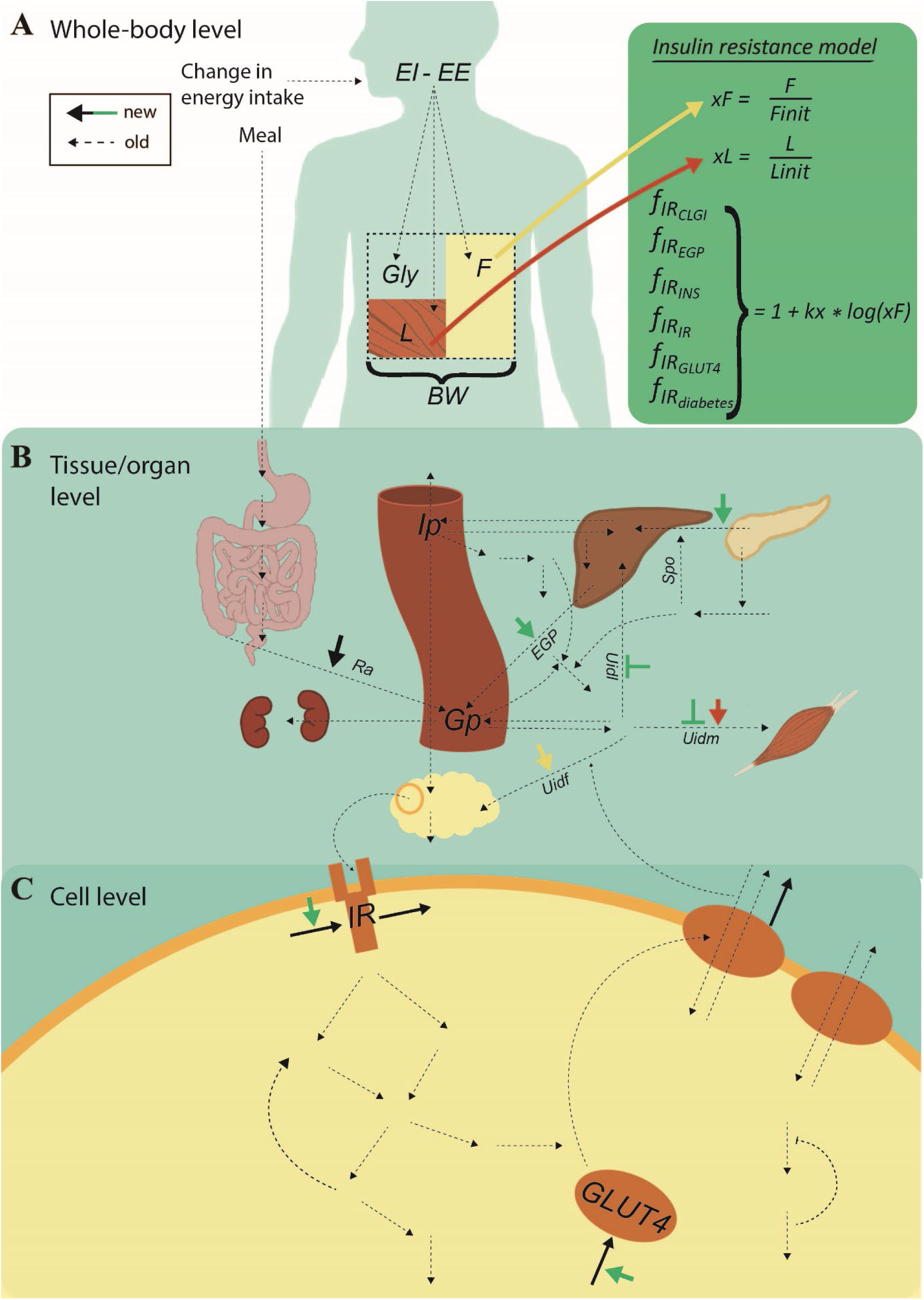
Detailed overview of the entire multi-level and multi-timescale model structure on the different levels. New reactions, added in this paper, are represented by solid lines, any color, while old reactions are represented with dashed lines. A) Whole-body level. The body composition model takes change in energy intake as input, i.e., the difference in energy intake (𝐸𝐼) and energy expenditure (𝐸𝐸). This difference translates to the outputs: changes in the masses of fat (𝐹), lean tissue (𝐿), and glycogen (𝐺𝑙𝑦). The total sum of these masses is the body weight (𝐵W). The insulin resistance model (green box) takes the change in fat mass (𝑥𝐹) as input. B) The following factors influence the glucose concentration on the tissue/organ level: the insulin resistance, 𝑥𝐹, the change in lean tissue (𝑥𝐿), and 𝐵W. More specifically, insulin resistance (green short arrows) increases endogenous glucose production (𝐸𝐺𝑃) and insulin secretion (𝐼𝑝𝑜), and decreases glucose uptake in both muscle (𝑈_𝑖𝑑𝑚_) and liver tissue (𝑈_𝑖𝑑𝑙_). Furthermore, 𝑥𝐿 increases 𝑈_𝑖𝑑𝑚_, 𝑥𝐹 increases glucose uptake in fat tissue (𝑈_𝑖𝑑_**_𝑓_**), and 𝐵W increases the rate of appearance of glucose (𝑅𝑎). C) Finally, the amount of insulin in fat tissue translates to insulin input on the cell level. More specifically, insulin binds to the insulin receptor (𝐼𝑅), causing a signaling cascade that ultimately results in glucose transporter 4 (𝐺𝐿𝑈𝑇4) being translocated to the plasma membrane to facilitate glucose transport. The new reactions on the cell level are the protein expressions of 𝐼𝑅 and 𝐺𝐿𝑈𝑇4 (black arrows going to), the effect of insulin resistance on the protein expression of 𝐼𝑅 and 𝐺𝐿𝑈𝑇4 (green arrows), as well as the degradation of 𝐼𝑅 and 𝐺𝐿𝑈𝑇4 (black arrows going out). These new reactions enable a gradual decrease in 𝐼𝑅 and 𝐺𝐿𝑈𝑇4, moving the cell towards diabetes.

### The model explains total weight change data and can correctly predicts data on all three levels in the Fast-food study

The whole-body model was trained on total body weight data, describing change in total body weight, obtained from a weight-increase study (23). In this study, the participants were told to increase their energy intake by around 3480 kcal per day by eating at least two extra meals of fast-food, and by decreasing their physical activity for four weeks (Fig. 3A). The model agrees well with the total body weight data, used for training the model (Fig. 3B). The model can also predict the increase in fat and fat free mass on the whole-body level (Fig. 3C). The interconnection between the whole-body and the tissue-level model was tested by comparing simulations from the entire multi-level model with tissue- and cell-level data from the weight-increase study (Fig. 3D-E). As can be seen in Fig. 3D, the experimental data for fasting insulin lies within the predicted bounds (light yellow area). The solid purple line shows the simulation with the lowest cost from the training to the weight data. Only one scaling parameter was adjusted to the data in Fig. 3D.

**Figure 3.**
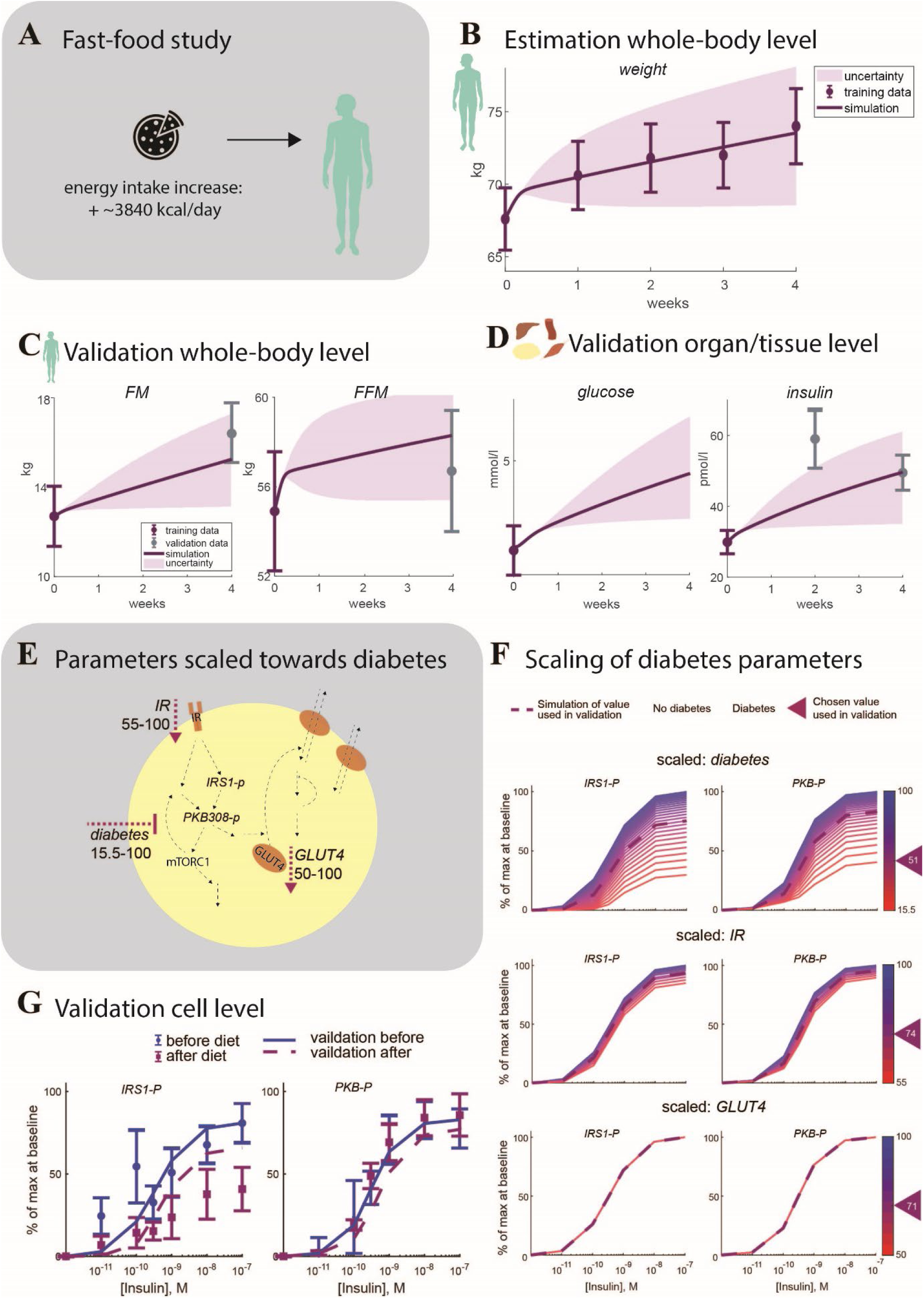
Results of model training and validation on Fast-food study data A). Comparison between model uncertainty (light purple area) for the best model simulation (the dark purple line) with the training data (purple error bars) or validation data (grey error bars). On the whole-body level, data for B) weight and C) fat mass (𝐹𝑀) and fat free mass (𝐹𝐹𝑀) was used for training and validation. On the tissue/organ level, data for D) glucose and insulin was used for training and validation. E) The diabetes effects on the cell level model – decrease in 𝐼𝑅, decrease in 𝐺𝐿𝑈𝑇4, and 𝑑𝑖𝑎𝑏𝑒𝑡𝑒𝑠, representing an attenuation of. F) Scaling of the three diabetes parameters (with the chosen values indicated with triangles) and the resulting behavior of the simulation curves as dose responses to insulin, to match the fit to data in G). Data and simulations of the dose responses of phosphorylated 𝑃𝐾𝐵, 𝑃𝐾𝐵308 − 𝑝 and phosphorylated 𝐼𝑅𝑆1, 𝐼𝑅𝑆1 − 𝑝 in response to the indicated concentrations of insulin for 10 min and normalized 0–100%. The predicted simulation before the Fast-food diet (blue solid line) use the non-diabetic parameters from (7) as they were, which gave a good agreement with data (blue error bars with circles). The three diabetes parameters were scaled to get the predicted simulation after the diet (purple dashed line) to fit to the corresponding data (purple error bars with squares).

The prediction of the cell-level insulin response data for the intracellular metabolites 𝐼𝑅𝑆1 − 𝑝 and 𝑃𝐾𝐵 − 𝑝 was scaled using two parameters to switch the three diabetes parameters (Fig. 3EF) in the model to 22% towards diabetes. As shown in Fig. 3G, this prediction also looks good, and is supported by a χ^2^ test, (𝑉(𝜃) = 19.3 < 21 = 𝜒^2^_𝑐𝑢𝑚,𝑖𝑛𝑣_(12,0.05)) for 𝐼𝑅𝑆1 − 𝑝 and (𝑉(𝜃) = 19.9 < 21 = 𝜒^2^_𝑐𝑢𝑚,𝑖𝑛𝑣_(12,0.05)) for 𝑃𝐾𝐵 − 𝑝.

### The multi-level model can predict whole-body-, tissue- and cell-level data based on weight increase data

All the simulations lie close to experimental data, as in Fig. 3B-D, G, meaning that the model can both explain training data and correctly predict independent validation data. It is therefore meaningful to look at predictions of other non-measured variables. The trained and validated model was therefore used to predict a continuation of the Fast-food diet for an additional 8 weeks, resulting in a continued weight increase (Fig. 4A left). During these additional weeks, the fasting plasma glucose and insulin levels reached prediabetic levels (Fig. 4A middle and right). The meal response of plasma and glucose also increased, while the glucose uptake in muscle and fat tissue decreased and increased respectively (Fig. 4B). The predictions of total 𝐼𝑅𝑆1 and 𝑃𝐾𝐵 expression at the cellular level got closer to the diabetic levels (Fig. 4D).

**Figure 4.**
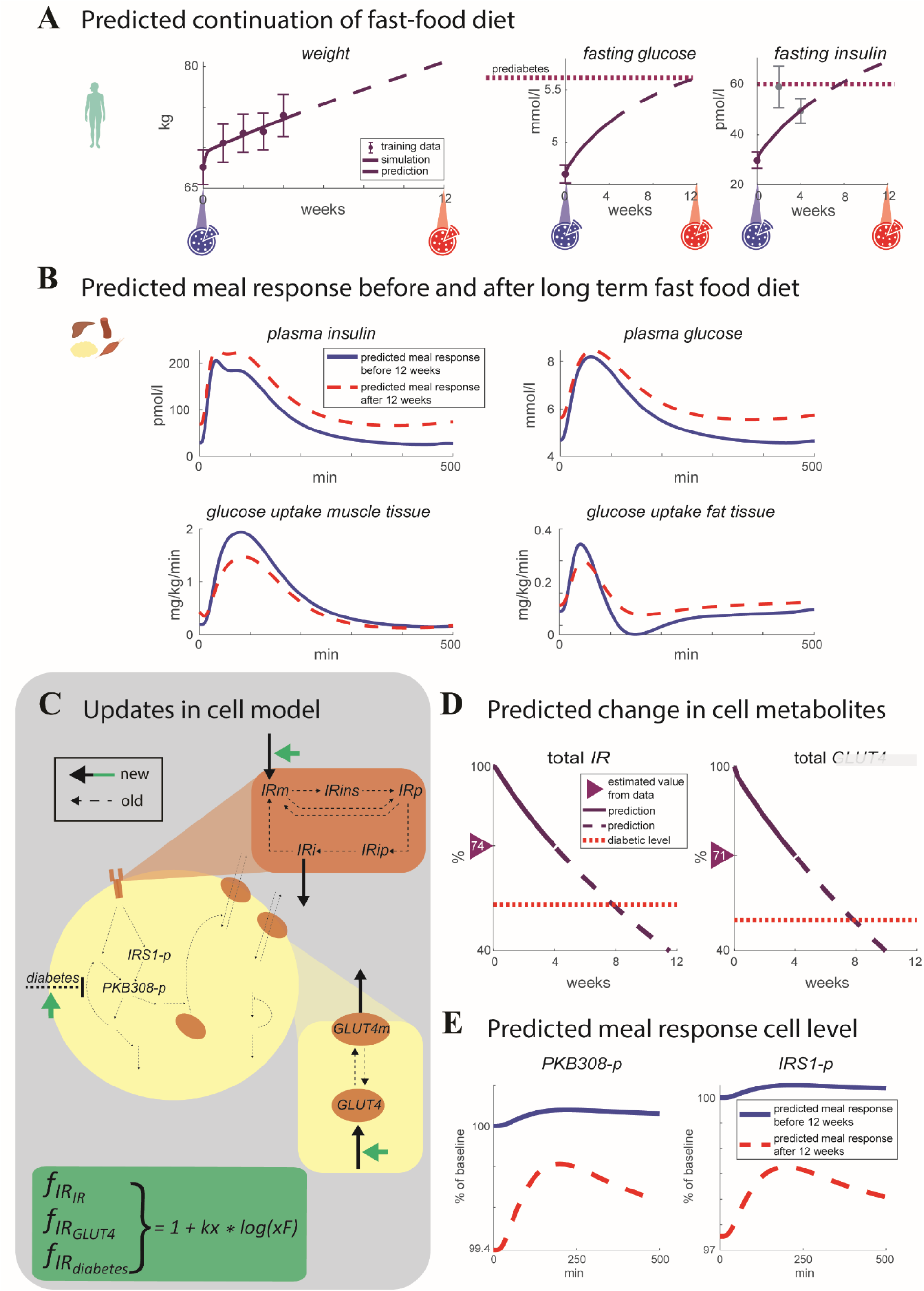
A) Model simulation of weight, fasting plasma glucose and insulin for a predicted continuation of the Fast-food diet for an additional 8 weeks. Prediabetic levels is shown as purple dotted line. Two meals are simulated during the period, before and the predicted 12 weeks (blue and red pizza icons respectively). B) Meal response simulations before (blue solid line) and after (red dashed line) the predicted 12-week Fast-food diet for plasma insulin, plasma glucose, and glucose uptake in muscle and fat tissue. C) The updates made to the cell level of the model and insulin resistance (green box). The added reactions include a protein expression 𝐼𝑅𝑚, degradation of 𝐼𝑅𝑖, protein expression of 𝐺𝐿𝑈𝑇4, and degradation of 𝐺𝐿𝑈𝑇4𝑚. The insulin resistance influences the protein expression of 𝐼𝑅𝑚 and 𝐺𝐿𝑈𝑇4 (green arrows). These updates enable the gradual change in total 𝐼𝑅 and 𝐺𝐿𝑈𝑇4 due to increased insulin resistance seen in D). After the 4 weeks of the Fast-food study, the total 𝐼𝑅 and 𝐺𝐿𝑈𝑇4 (solid purple) have reached the values estimated from data in Fig. 3FG. After the additional 8 weeks, total 𝐼𝑅 and 𝐺𝐿𝑈𝑇4 (dashed purple) have gone down further towards but not completely reached the diabetic value (dotted red line). E) Cell response to the simulated meals before (blue solid line) and after the 12 weeks (red dashed line), specifically the response of 𝑃𝐾𝐵308 − 𝑝 and 𝐼𝑅𝑆1 − 𝑝.

### The model describes and predicts weight changes from Topiramate study

The model was further validated on a weight-decrease study with the drug topiramate (22). The model was trained on two doses of topiramate - 64 and 192 mg/day - and then validated on a third dosage - 96 mg/dl. The model training passes a χ^2^ test (𝑉(𝜃) = 3.0 < 36 = 𝜒^2^_𝑐𝑢𝑚,𝑖𝑛𝑣_(24,0.05)). As shown in Fig. 5, the validation lies within the predicted bounds, and it also passes a χ^2^ test (𝑉(𝜃) = 4.8 < 21 = 𝜒^2^_𝑐𝑢𝑚,𝑖𝑛𝑣_(12,0.05)).

**Figure 5.**
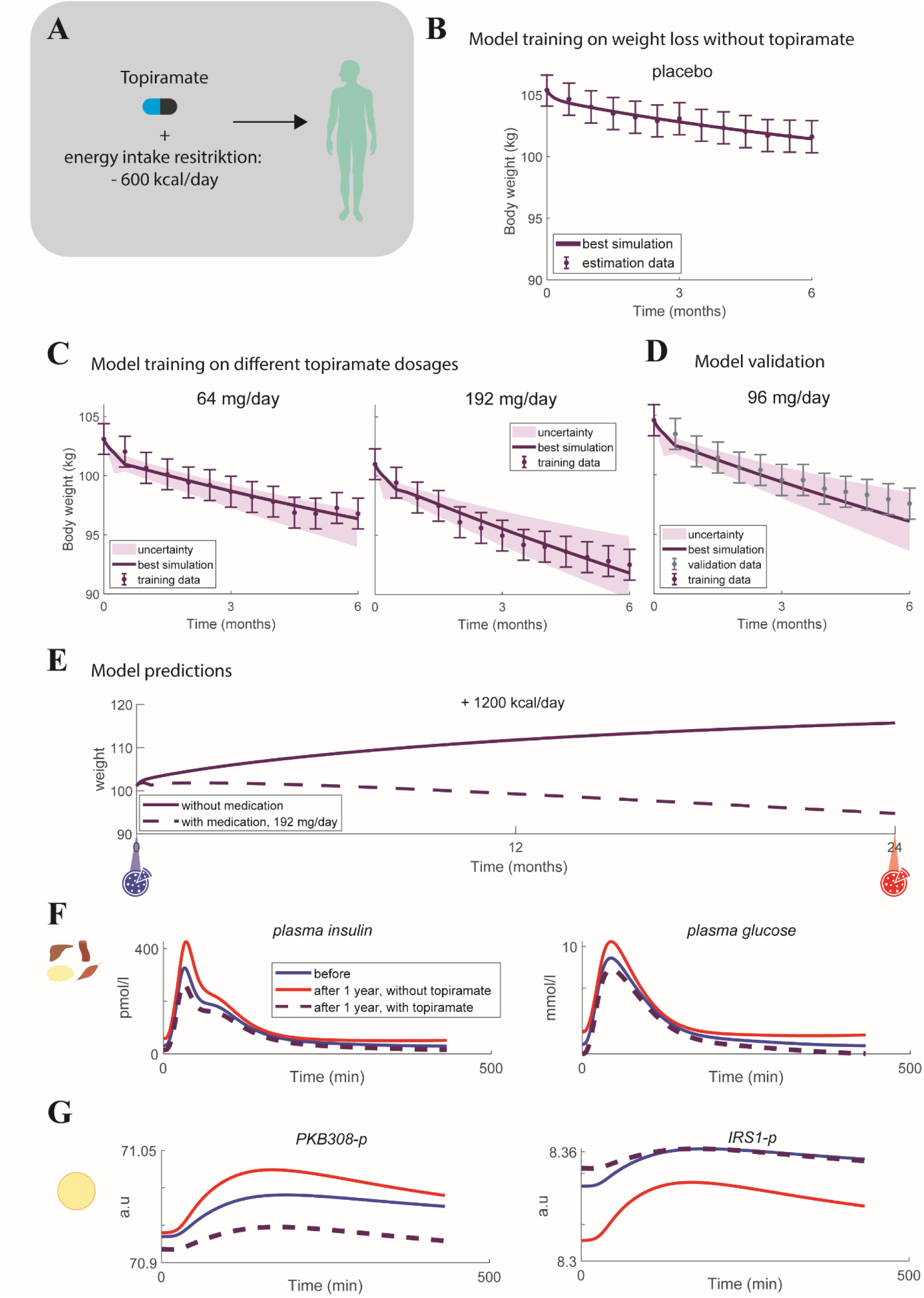
A) Overview of the Topiramate study, in which the patients were treated with three different dosages of the weight-loss drug topiramate, and instructed to eat on average 600 kcal less per day and also took different dosages of the weight loss drug Topiramate. The results of the model training and validation on topiramate data are shown as purple lines for the best simulation, purple error bars for the data, and shaded purple areas for model uncertainty, for the B) fit to placebo data, C) fit to weight data for Topiramate dosages 64 mg/day and 192 mg/day, and D) model validation on Topiramate dosage 96 mg/day, where the validation data is shown as gray error bars. The first data point was used to set initial conditions for the corresponding simulations. E) The trained model was used to make predictions made for two different scenarios not done during the Topiramate study: increasing the energy intake with 1200 kcal/day for 1 year, without topiramate treatment (solid purple line) and with 192 mg/day topiramate treatment (dashed purple line). F) Predictions of meal responses before the predicted diet and topiramate intervention (blue solid line), after 1 year of energy intake increase without treatment (red solid line) and with treatment (purple dashed line) for 𝑝𝑙𝑎𝑠𝑚𝑎 𝑖𝑛𝑠𝑢𝑙𝑖𝑛 and 𝑝𝑙𝑎𝑠𝑚𝑎 𝑔𝑙𝑢𝑐𝑜𝑠𝑒 on the organ/tissue level, and G) 𝐼𝑅𝑆1 − 𝑝 and 𝑃𝐾𝐵308 − 𝑝 on the cell level.

### The multi-level model can predict tissue- and cell-level data based on weight decrease

As shown in Fig. 5BCD, the simulations describe accurately both experimental estimation and validation data. It is therefore, as with Fig. 3BCDG and the weight-increase scenario, meaningful to look at predictions for the population in the Topiramate study as well. Such predictions on the organ/tissue- and cell level were made using the fit of the whole-body model to the weight data from the Topiramate study. Specifically, two scenarios that were not part of the Topiramate study were both predicted and compared: 1) an increase in energy intake by 1200 kcal per day for 1 year without topiramate treatment, and 2) the same increase in energy intake (1200 kcal/day for 1 year) but with topiramate treatment, 192 mg/day (Fig. 5E). In the first scenario, the weight increases with almost 15 kg (Fig. 5E, solid line), while in the scenario with topiramate, the model predicts a decrease in weight (Fig. 5E, dashed line), despite the increase in calories. When looking at a meal response at the organ/tissue level, before and after the predicted year of weight increase or decrease (Fig. 5F), both the plasma insulin and glucose levels have increased after one year without drug treatment (red solid lines) compared with before (blue solid lines). After one year of energy-intake increase with topiramate treatment (purple dashed lines), the plasma insulin and glucose levels have instead decreased slightly. Similar changes can also be seen in the meal response on the cellular level (Fig. 5G) – 𝑃𝐾𝐵308 − 𝑝 protein levels have increased after 1 year of only increase in energy intake compared to before, and the same protein level had decreased after one year of topiramate treatment, while 𝐼𝑅𝑆1 − 𝑝 has decreased after 1 year of energy intake increase only and slightly decreased after 1 year on topiramate.

### Outline of possible future clinical usage

Predictions such as the ones made in Fig. 4 and Fig. 5E-G can, among other things, potentially be used in health care. More specifically, an active version of a digital twin, which has been personalized using data from one person (Fig. 6A), can be used to simulate and predict different scenarios (Fig. 6B). For example, it is possible simulate how different diets can result in either an increase or decrease in weight, such as the two scenarios shown in Fig. 6B. Such simulated scenarios can then be compared with each other, either for pedagogical and motvational purposes or for treatment evaluation. When used for pedagogical and motivational purposes, the simulations can be used to increase the understanding of the physiological effects that different lifestyles and/or treatments have on your physiology over extended periods of time. Such an increased understanding could then hopefully lead to better motivation to follow a certain lifestyle or treatment intervention. When using the simulations for treatment evaluation, the scenarios can be compared in order to chose the lifestyle and/or treatment most suited for the particular person using it, both in terms of outcome (e.g. which diet results in the most decreased risk of diabetes) and what changes you can and are willing to do in your life (e.g. which diet with a good enough outcome could you see yourself comply to). Finally, the multi-time scale and multi-level aspect of the model can potentially be utilized to get continuous feedback on the chosen lifestyle and/or treatment (Fig. 6C), by zooming in on shorter time scales (as in Fig. 6C big blue box) and comparing with collected data, or simulating something else in the digital twin (e.g. the glucose response in plasma following a meal, as in Fig. 6C left and right small boxes). This feedback can help to evaluate the life style – does this chosen intervention seem to work for me as predicted?

**Figure 6:**
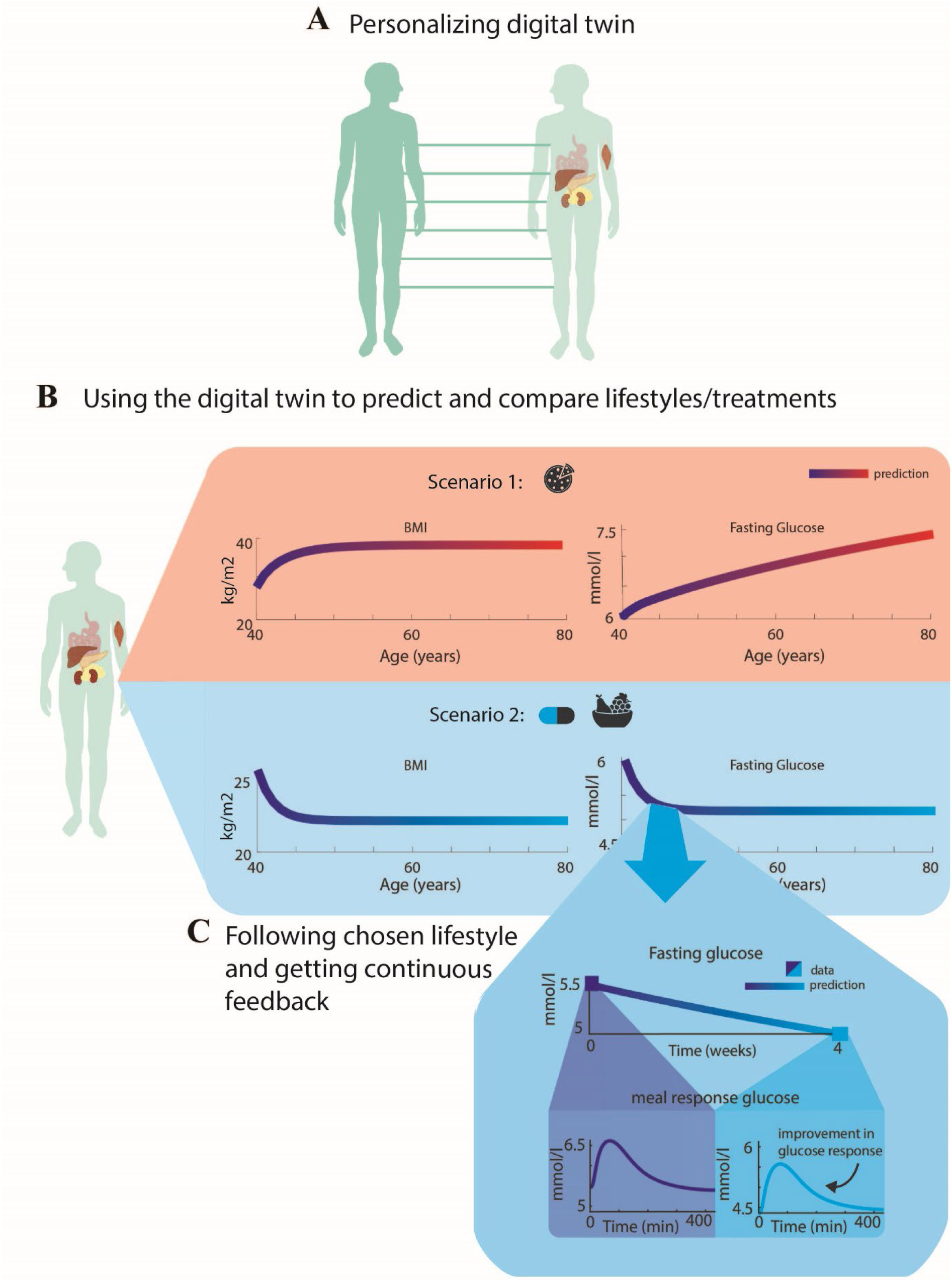
A) Personalizing a digital twin using data from one person to train and validate a passive digital twin, such as the one presented herein, and making the digital twin active. B) Using the digital twin to predict and compare scenarios with different lifestyles and/or treatments. In this example, the digital twin is used to predict two scenarios. In scenario 1, the digital twin simulates an increase in energy intake for 40 years (from 40 to 80 years of age) and a resulting increase in BMI – from overweight to obese levels (BMI over 25 and 30 kg/m^2^, respectively) - and an increase in fasting plasma glucose – from prediabetic to diabetic levels (fasting glucose above 5.6 and 7 mmol/l, respectively). In Scenario 2, the digital twin simulates a decrease in energy intake with a weight-loss drug such as topiramate, resulting in a decrease to healthy levels of BMI and fasting plasma glucose. C) Following the chosen lifestyle and getting continuous feedback by zooming in on 4 weeks of the predicted fasting plasma glucose (solid line) and comparing with data (blue squares) collected by the user. Zooming in even more and looking at meal response glucose before and after the 4 weeks, one can see that the glucose curve is higher before (left box) compared to after (right box), indicating an improvement in meal response glucose levels as well.

## Discussion

### Summary of main findings

Herein, we have presented a first multi-level, multi-timescale, and mechanistic model of the progression of insulin resistance in humans. The model describes insulin resistance development on three different biological levels: whole-body composition (Fig. 2A, 3BC, 4A, and 5C), plasma glucose and insulin (Fig. 2B, 3D, 4B, and 5D), and intracellular adipocyte insulin signaling (Fig. 2C, 3G, 4DE and 5E). The model agrees with the multi-level dataset from the Fast-food study (17,18), both describing estimation data (Fig. 3B) and correctly predicting independent validation data (Fig. 3CDG). For this weight-increase study, we predict traditional biomarkers which had not been measured, such as oral glucose tolerance test and other intracellular signaling intermediaries (Fig. 4). The model also agrees with whole-body weight-loss data from the Topiramate study (Fig. 5BC) (22). Moreover, for this study, we use the model to predict changes in a glucose tolerance test and intracellular insulin signaling (Fig. 5DE). Finally, we illustrate how this model potentially can be used to improve health in future eHealth technologies (Fig. 6).

### New strengths and possibilities with our new modular and multi-scale model for insulin resistance

An important strength of this model is that it combines three well-determined and validated models into an interconnected multi-scale model. Having a multi-scale model is an important strength since the progression of diabetes in reality is multi-scale, as seen in the data (Fig. 3). Despite this importance, there existed no previously available multi-scale model that could describe such data. Nevertheless, there exists models that describe the different levels separately: the Hall model for whole-body weight describes changes over months and years (3); the Dalla Man model for the meal response describes the interplay between plasma glucose and insulin (4); and the Brännmark model that describes intracellular insulin signaling data in adipocytes (6,7). However, these three models had previously not been connected into a single model, in part because the arguably most central connection between them – adiposity-driven insulin resistance – had not previously been modelled. Herein, we have for the first time connected these three well-established models and levels into a multi-scale model, by introducing a new model for the progression of insulin resistance. This is the first such human, multi-scale insulin resistance model. The connecting kit, the adiposity-driven insulin resistance model, has been adopted from a corresponding multi-scale model for mice (11), even though the three constituent models for the three levels and timescales, come from existing models that were specific to humans. Moreover, the cell level model has also been adjusted to allow a continuous development of insulin resistance, by allowing some of the model’s steady states to instead change over time. A final important aspect of this multi-scale model is that it is modular, meaning that the different subsystems and organs described in the model can be changed to other models with more or less details (25,26).

Another strength with our new model is that it can describe not only estimation data, but also correctly predict independent validation data, and can thus also be used to predict non-measured variables. The model correctly describes estimation data of weight change from both the Fast-food study (Fig. 3B) and the Topiramate study (Fig. 5C). Furthermore, the model also correctly described independent validation data from both these studies. For the Fast-food study, the model describes independent data on changes in fat mass and fat free mass (Fig. 3C), fasting glucose and insulin concentrations (Fig. 3D), and the insulin response of the intracellular signaling metabolites (Fig. 3E). In all these predictions, the model only changed one parameter: the scale difference between mice and humans in the insulin resistance model. For the Topiramate study, the model describes independent data for weight change using a dosage of topiramate that was not used for fitting, 96 mg/day (Fig. 5D). Because of the success of these validation tests, we then used the model to make predictions of the gradual changes of some of the things that were not measured in the original study. For example, we could predict how the glucose levels and fluxes changes during the studies, as well as how intracellular signaling is changing. These kinds of predictions are something that the earlier model cannot do, since they require the interplay between the different layers. These predictions of additional non-measured variables can in principle be tested by doing new studies where these variables are measured, and this could either validate the current model even further, or reject the model, and both these outcomes would provide new mechanistic insights regarding the progression of insulin resistance.

### Limitations with our model

The current version of the model has some limitations. One such limitation is that the implementation of the fat-dependent insulin resistance is a minimal model, using simple relatively expressions. Specifically, the model lacks relevant details and hypotheses assumed to be involved in insulin progression. One such mechanistic hypothesis is ectopic fat storage and inflammation in liver and pancreas (27,28). Inflammation is also often believed to play a role in the adipose tissue itself, as is the varying cell size distributions of adipocytes (29,30). These things could be included in future, more detailed versions, of the model. However, all of these are processes that are not covered by the model’s current level of detail, and among processes currently included, the progression of insulin resistance is mechanistic, in the sense that it affects the right included mechanisms. For instance, the EGP-production of glucose from the liver is known to be impacted by insulin resistance, and this impact is included, even though the underlying mechanisms for this impact are not included. To include such underlying mechanisms would allow us to simulate a wider array of drugs, including e.g. anti-inflammatory drugs like cd44-inhibitors (31,32), or drugs that influence the size of adipocytes like metformin (33). These potential additions could thus be useful for both drug development and individualized prevention. Apart from this lack of mechanistic detail, the current implementation of insulin resistance progression is given by a logarithmic expression (Eq. 3-5,10-12). This expression thus excludes potential transient and/or adjustment processes in the body. Also, the current progression of insulin resistance has only been validated on a relatively small weight span and population, meaning that higher or lower weight changes and other time scales might not be accurately represented by the model.

Another potential limitation with the current model concerns how the interconnection was introduced. Specifically, the interconnection is top down only – the whole-body level only influences the organ/tissue level only goes in one direction, that is from the top-level (whole-body) to the lower level (organ/tissue/cell), and is not reversible. This implementation of the connection means that the meal response or meal response dynamics does not affect the whole-body composition changes, which, in reality, it does. A future implementation of the interconnection could describe how short-term changes in meal response dynamics would lead to short-term changes in ectopic fat storage, which over time would lead to long-term changes in fat mass, and therefore also overall body weight. To implement such a two-way interconnection between the levels, the model should represent fat tissue in greater detail, including e.g., proliferation and death of adipocytes, the effects of differently sized adipocytes, the amount of fat in each adipocyte, and ectopic fat storage. (29,30,34,35). Other more realistic interconnections include for example different hunger and fat-mass regulating hormones (such as leptin, adiponectin, various inflammation mechanisms, intracellular mechanisms on more organs than fat tissue), as well as the interplay between glucose, proteins, and fat (20,36–39).

A third potential drawback is that the model is heavily focused on adipose tissue and the adipocytes, and their involvement in insulin resistance. This adipocentric explanation is one of the most popular ones for the progression of insulin resistance, but not the only one. Other explanations do exist, such as various genetic explanations, inflammation in other organs and/or due to ectopic fat storage (29,29,34,35,40–42). It is also possible that there are several mechanisms leading to insulin progression that are true at the same time or for different clusters of people with insulin resistance. There are also at least some evidence for the existence of a range of different diabetic subtypes (43,44), and that there are also different possible pathways to diabetes and insulin resistance (9). Ideally, all different hypotheses should be implemented and compared.

### Future applications of our multi-scale model: digital twins, eHealth, and drug development

The multi-scale model presented herein is a so called passive digital twin. A passive digital twin is, in contrast to an active digital twin, not personalized using individual data, even though it could be. Both active and passive digital twins can be useful in an eHealth scenario. Passive twins can for example be used to describe general dynamics of disease progression and be used as a medical pedagogics tool. For example, when looking at the progression of insulin resistance, the model can show how an increased energy intake can result in a weight increase, and eventually also to progression towards insulin resistance and type 2 diabetes. To simulate such illustrations of the effect of daily habits could both help to convey medical knowledge in a comprehensive way and motivate to making life-style changes. Active digital twins can also help with medical pedagogics and motivation, but with the additional benefit of being able to make personalized predictions. Such predictions could potentially also be used to help motivate patients to adhere to prescribed drugs or to more stringently follow their prescribed diet and exercise-schemes. Furthermore, mechanistically based, multi-scale models for the progression of insulin resistance and type 2 diabetes could potentially also be used to evaluate different care interventions. For example, when using weight loss as a prevention or treatment for diabetes, a digital twin can be used for comparison of different options - topiramate could be compared to other interventions, both by comparing the effects on weight loss and other relevant biomarkers. All of these potential applications of a digital twin could be further increased by connecting the digital twin with a machine learning risk model-based drug development, and systems pharmacology, creating a hybrid model. This hybrid model could then be used to calculate a personalized or general risk for different diseases, like diabetes or cardiovascular diseases, given a certain scenario simulated by the digital twin. Then, when comparing different weight-loss drugs, their relative effect on the risk of disease could also be compared (45,46). In conclusion, the multi-scale model presented herein constitutes the basis for an active or passive digital twin technology that could be used to aid medical pedagogics and increase motivation and compliance, and can as such aid in prevention and treatment of insulin resistance.

## Acknowledgements

We thank MSc Maria Kjellsson for her valuable input on the manuscript.

