## Supplementary for "A multi-scale digital twin for adiposity-driven insulin resistance in humans: diet and drug effects"

All the ODEs for the final model:

$$\frac{d}{dt}(IRm) = -v1a - v1basal + v1r + v1g + vIR \tag{1}$$
$$\frac{d}{dt}(IRp) = v1basal + v1c - v1d - v1g \tag{2}$$
$$\frac{d}{dt}(IRins) = v1a - v1c \tag{3}$$
$$\frac{d}{dt}(IRip) = v1d - v1e \tag{4}$$
$$\frac{d}{dt}(IRi) = v1e - v1r - vIRdeg \tag{5}$$
$$\frac{d}{dt}(IRS1) = v2b + v2g - v2a - v2basal \tag{6}$$
$$\frac{d}{dt}(IRS1p) = v2a + v2d - v2b - v2c \tag{7}$$
$$\frac{d}{dt}(IRS1p307) = v2c - v2d - v2f \tag{8}$$
$$\frac{d}{dt}(IRS1307) = v2basal + v2f - v2g \tag{9}$$
$$\frac{d}{dt}(X) = v3b - v3a \tag{10}$$
$$\frac{d}{dt}(Xp) = v3a - v3b \tag{11}$$
$$\frac{d}{dt}(PKB) = -v4a + v4b + v4h \tag{12}$$
$$\frac{d}{dt}(PKB308p) = v4a - v4b - v4c \tag{13}$$
$$\frac{d}{dt}(PKB473p) = -v4e + v4f - v4h \tag{14}$$
$$\frac{d}{dt}(PKB308p473p) = v4c + v4e - v4f \tag{15}$$
$$\frac{d}{dt}(mTORC1) = v5b - v5a \tag{16}$$
$$\frac{d}{dt}(mTORC1a) = v5a - v5b \tag{17}$$
$$\frac{d}{dt}(mTORC2) = -v5c + v5d \tag{18}$$
$$\frac{d}{dt}(mTORC2a) = v5c - v5d \tag{19}$$
$$\frac{d}{dt}(AS160) = v6b1 - v6f1 \tag{20}$$
$$\frac{d}{dt}(AS160p) = v6f1 - v6b1 \tag{21}$$

$$\frac{d}{dt}(GLUT4m) = (v7f - v7b - vGLUTdeg) \tag{22}$$
$$\frac{d}{dt}(GLUT4) = -v7f + v7b + vGLUT \tag{23}$$
$$\frac{d}{dt}(S6K) = v9b1 - v9f1 \tag{24}$$
$$\frac{d}{dt}(S6Kp) = v9f1 - v9b1 \tag{25}$$
$$\frac{d}{dt}(S6) = v9b2 - v9f2 \tag{26}$$
$$\frac{d}{dt}(S6p) = v9f2 - v9b2 \tag{27}$$
$$\frac{d}{dt}(G_p) = EGP + Ra - E - U_{ii} - k_1 \cdot G_p + k_2 \cdot G_t \tag{28}$$
$$\frac{d}{dt}(G_t) = -U_{id} + k_1 \cdot G_p - k_2 \cdot G_t - U_{idl} \tag{29}$$
$$\frac{d}{dt}(I_l) = (-m_1 \cdot I_l) - m_3 \cdot I_l + m_2 \cdot I_p + S \tag{30}$$
$$\frac{d}{dt}(I_p) = (-m_2 \cdot I_p) - m_4 \cdot I_p + m_1 \cdot I_l \tag{31}$$
$$\frac{d}{dt}(Q_{sto1}) = -k_{gri} \cdot Q_{sto1} \tag{32}$$
$$\frac{d}{dt}(Q_{sto2}) = (-k_{empt} \cdot Q_{sto2}) + k_{gri} \cdot Q_{sto1} \tag{33}$$
$$\frac{d}{dt}(Q_{gut}) = (-k_{abs} \cdot Q_{gut}) + k_{empt} \cdot Q_{sto2} \tag{34}$$
$$\frac{d}{dt}(I_1) = -k_i \cdot (I_1 - I) \tag{35}$$
$$\frac{d}{dt}(I_d) = -k_i \cdot (I_d - I_1) \tag{36}$$
$$\frac{d}{dt}(INS_f) = (-p_{2U} \cdot INS_f) + p_{2U} \cdot (I - I_b) \tag{37}$$
$$\frac{d}{dt}(I_{po}) = ((-gamma \cdot I_{po}) + S_{po}) \cdot f_{IR_{Ins}} \tag{38}$$
$$\frac{d}{dt}(Y) = -alpha \cdot (Y - beta \cdot (G - G_b)) \tag{39}$$
$$\frac{d}{dt}(INS) = V_1 - V_2 \tag{40}$$
$$\frac{d}{dt}(Glu_{in}) = p1 \cdot (V_{in} - V_{out}) - V_{G6P} \tag{41}$$
$$\frac{d}{dt}(G6P) = V_{G6P} - V_{met} \tag{42}$$

$$\frac{d}{dt}(Gly) = \frac{v1}{rhog} \tag{43}$$

$$\frac{d}{dt}(ECF) = v2 \tag{44}$$

$$\frac{d}{dt}(F) = \frac{(1 - v3) \cdot (k1 - v1)}{rhof} \tag{45}$$

$$\frac{d}{dt}(L) = \frac{v3 \cdot (k1 - v1)}{rhol} \tag{46}$$

$$\frac{d}{dt}(AT) = v4 \tag{47}$$

$$\frac{d}{dt}(A) = -Ka \cdot A \tag{48}$$

$$\frac{d}{dt}(C) = Ka \cdot \frac{A}{V} - K23 \cdot C + K23 \cdot y3 - k10 \cdot C \tag{49}$$

$$\frac{d}{dt}(C1) = K32 \cdot C - K32 \cdot C1 \tag{50}$$

All variables of final model:

$$aa = \frac{5/2}{(1-b)/D} \tag{51}$$

$$cc = \frac{5/2}{d/D} \tag{52}$$

$$EGP = (k_{p1} - k_{p2} \cdot G_p - k_{p3} \cdot I_d - k_{p4} \cdot I_{po}) \cdot f_{IR_{EGP}} \tag{53}$$

$$V_{lmax} = V_l + V_{lX} \cdot INS \tag{54}$$

$$V_{mmax} = V_m + V_{mX} \cdot INS \tag{55}$$

$$E = 0 \tag{56}$$

$$S = gamma \cdot I_{po} \tag{57}$$

$$I = \frac{I_p}{V_I} \tag{58}$$

$$G = \frac{G_p}{V_G} \tag{59}$$

$$HE = (-m_5 \cdot S) + m_6 \tag{60}$$

$$m_3 = HE \cdot \frac{m_1}{(1-HE)} \tag{61}$$

$$Q_{sto} = Q_{sto1} + Q_{sto2} \tag{62}$$

$$Ra = f \cdot k_{abs} \cdot \frac{Q_{gut}}{BW} \tag{63}$$

$$k_{empt} = k_{min} + \frac{(k_{max} - k_{min})}{2} \cdot (tanh(aa \cdot (Q_{sto} - b \cdot D)) - tanh(cc \cdot (Q_{sto} - d \cdot D)) + 2) \tag{64}$$

$$bf = (be + kbf \cdot (INS_f + INS_{\text{offset}})) \cdot bradykinin \tag{65}$$

$$bfe_f = (bf - bf_b) \cdot (INS_f - INS_b) \cdot p_{bf} \tag{66}$$

$$S_{po} = Y + K \cdot \frac{(EGP + Ra - E - U_{ii} - k_1 \cdot G_p + k_2 \cdot G_t)}{V_G} + S_b \tag{67}$$

$$INS_{fe} = nC \cdot (k8 \cdot \frac{GLUT4m}{pf} + \frac{GLUT1}{pf} + bfe_f) \tag{68}$$

$$V_{in} = p4 \cdot G_t \cdot INS_{fe} \tag{69}$$

$$V_{out} = p3 \cdot Glu_{in} \tag{70}$$

$$U_{idm} = xLV_{mmax} \cdot \frac{G_t}{(K_m + G_t) f_{IR_{CLGI}}} \tag{71}$$

$$U_{idl} = V_{lmax} \cdot \frac{G_t}{(K_l + G_t) f_{IR_{CLGI}}} \tag{72}$$

$$U_{idf} = xFp_5 \cdot (V_{in} - V_{out}) \tag{73}$$

$$U_{id} = U_{idf} + U_{idm} + U_{idl} \tag{74}$$

$$U = U_{ii} + U_{id} + U_{idl} \tag{75}$$

$$V_2 = k_2 \cdot INS \tag{76}$$

$$V_1 = k_1 \cdot (I - I_b) \tag{77}$$

$$v1a = IRm \cdot k1a \cdot (INS_f + 5) \cdot 1e - 3 \tag{78}$$
$$v1basal = k1basal \cdot IRm \tag{79}$$
$$v1c = IRins \cdot k1c \tag{80}$$
$$v1d = IRp \cdot k1d \tag{81}$$
$$v1e = IRip \cdot k1f \cdot Xp \tag{82}$$
$$v1g = IRp \cdot k1g \tag{83}$$
$$v1r = IRi \cdot k1r \tag{84}$$
$$v2a = IRS1 \cdot k2a \cdot IRip \tag{85}$$
$$v2b = IRS1p \cdot k2b \tag{86}$$
$$v2c = IRS1p \cdot k2c \cdot mTORC1a \cdot diabetes \tag{87}$$
$$v2d = IRS1p307 \cdot k2d \tag{88}$$
$$v2f = IRS1p307 \cdot k2f \tag{89}$$
$$v2basal = IRS1 \cdot k2basal \tag{90}$$
$$v2g = IRS1307 \cdot k2g \tag{91}$$
$$v3a = X \cdot k3a \cdot IRS1p \tag{92}$$
$$v3b = Xp \cdot k3b \tag{93}$$
$$v5a = mTORC1 \cdot (k5a1 \cdot PKB308p473p + k5a2 \cdot PKB308p) \tag{94}$$
$$v5b = mTORC1a \cdot k5b \tag{95}$$
$$v5c = mTORC2 \cdot k5c \cdot IRip \tag{96}$$
$$v5d = k5d \cdot mTORC2a \tag{97}$$
$$v4a = k4a \cdot PKB \cdot IRS1p \tag{98}$$
$$v4b = k4b \cdot PKB308p \tag{99}$$
$$v4c = k4c \cdot PKB308p \cdot mTORC2a \tag{100}$$
$$v4e = k4e \cdot PKB473p \cdot IRS1p307 \tag{101}$$
$$v4f = k4f \cdot PKB308p473p \tag{102}$$
$$v4h = k4h \cdot PKB473p \tag{103}$$
$$v6f1 = AS160 \cdot (k6f1 \cdot PKB308p473p + k6f2 \cdot \frac{PKB473p^n6}{(km6^n6 + PKB473p^n6))} \tag{104}$$
$$v6b1 = AS160p \cdot k6b \tag{105}$$
$$v7f = GLUT4 \cdot k7f \cdot AS160p \tag{106}$$
$$v7b = GLUT4m \cdot k7b \tag{107}$$
$$v9f1 = S6K \cdot k9f1 \cdot \frac{mTORC1a^n9}{km9^n9 + mTORC1a^n9} \tag{108}$$
$$v9b1 = S6Kp \cdot k9b1 \tag{109}$$
$$v9f2 = S6 \cdot k9f2 \cdot S6Kp \tag{110}$$
$$v9b2 = S6p \cdot k9b2 \tag{111}$$
$$V_{G6P} = V_{G6P_{max}} \cdot \frac{Glu_{in}}{(k_{gluin} + Glu_{in})} \cdot \frac{1}{(k_{G6P} + G6P)} \tag{112}$$
$$V_{met} = p3 \cdot G6P \tag{113}$$

$$xF = \frac{F}{Finit} \tag{114}$$

$$xL = \frac{L}{Linit} \tag{115}$$

$$f_{IR_{CLGI}} = 1 + (b_{CLGI} \cdot log(xF)) \cdot scale \tag{116}$$

$$f_{IR_{EGP}} = 1 + (b_{EGP} \cdot log(xF)) \cdot scale \tag{117}$$

$$f_{IR_{Ins}} = 1 + (b_{Ins} \cdot log(xF)) \cdot scale \tag{118}$$

$$f_{IR_{ir}} = 1 - b_{IR} \cdot log(xF) \tag{119}$$

$$f_{IR_{glut4}} = 1 - b_{GLUT} \cdot log(xF) \tag{120}$$

$$diabetes = 1 - b_{diabetes} \cdot log(xF) \tag{121}$$

$$CC = 10.4 \cdot \frac{rhol}{rhof} \tag{122}$$

$$p = \frac{CC}{(CC + F)} \tag{123}$$

$$EEinit = PAE \cdot RMRinit \tag{124}$$

$$EIninit = EEinit \tag{125}$$

$$CIninit = fCIn \cdot EIninit \tag{126}$$

$$kg = \frac{CIninit}{Ginit \cdot Ginit} \tag{127}$$

$$dEI_init = EIrestriction \cdot joules \tag{128}$$

$$IC = 1 - lmax \cdot \frac{C^{h2}}{(C^{h2} + IC_{50}^{h2})} \tag{129}$$

$$EI_{vehicle} = EIninit + dEI_{init} - (dEI_{init} - dEI_{ss} \cdot joules) \cdot \frac{time^{h1}}{(time^{h1} + t_{half}^{h1})} \tag{130}$$

$$EIn = EI_{vehicle} \cdot IC \tag{131}$$

$$CInvalue = fCIn \cdot EI_{vehicle} \tag{132}$$

$$CIn = CIninit \cdot CInvalue \tag{133}$$

$$PAL = PAE \tag{134}$$

$$BW = (F + L + (1 + 2.7) \cdot Gly + ECF) \tag{135}$$

$$TEF = bTEF \cdot (EIn - EIninit) \tag{136}$$

$$KK = EEinit - (gf \cdot Finit + gl \cdot Linit + delta \cdot BWinit) \tag{137}$$

$$EE1 = -BW \cdot delta \cdot rhof \cdot rhol - KK \cdot rhof \cdot rhol - rhof \cdot rhol \cdot TEF - rhof \cdot rhol \cdot AT \quad (138)$$

$$+ etal \cdot p \cdot rhof \cdot CIn + etaf \cdot rhol \cdot CIn - etaf \cdot p \cdot rhol \cdot CIn - etal \cdot p \cdot rhof \cdot EIn \quad (139)$$

$$- etaf \cdot rhol \cdot EIn + etaf \cdot p \cdot rhol \cdot EIn - gf \cdot rhof \cdot rhol \cdot F \quad (140)$$

$$- etal \cdot kg \cdot p \cdot rhof \cdot Gly \cdot Gly - etaf \cdot kg \cdot rhol \cdot Gly \cdot Gly \quad (141)$$

$$+ etaf \cdot kg \cdot p \cdot rhol \cdot Gly \cdot Gly - gl \cdot rhof \cdot rhol \cdot L \quad (142)$$

$$EE = \frac{EE1}{(-etal \cdot p \cdot rhof - etaf \cdot rhol + etaf \cdot p \cdot rhol - rhof \cdot rhol)} \quad (143)$$

$$k1 = EIn - EE \quad (144)$$

$$vIR = kIR \cdot f_{IR_{ir}} \quad (145)$$

$$vIRdeg = kIRdeg \cdot IRi \quad (146)$$

$$vGLUT = kGLUT \cdot f_{IR_{glut4}} \quad (147)$$

$$vGLUTdeg = kGLUTdeg \cdot GLUT4m \quad (148)$$

$$IRtot = IRm + IRins + IRp + IRip + IRi \quad (149)$$

$$GLUT4tot = GLUT4 + GLUT4m \quad (150)$$

| Annotation | Value | Annotation | Value |
|---|---|---|---|
| $k1a$ | 0.633141 | $k6f2$ | 36.9348 |
| $k1basal$ | 0.0331338 | $km6$ | 30.5424 |
| $k1c$ | 0.876805 | $n6$ | 2.13707 |
| $k1d$ | 31.012 | $k6b$ | 65.1841 |
| $k1f$ | 1839.58 | $k7f$ | 50.9829 |
| $k1g$ | 1944.11 | $k7b$ | 2285.97 |
| $k1r$ | 0.547061 | $k8$ | 724.242 |
| $k2a$ | 3.22728 | $glut1$ | 7042.19 |
| $k2c$ | 5758.78 | $k9f1$ | 0.12981 |
| $k2basal$ | 0.0422768 | $k9b1$ | 0.0444092 |
| $k2b$ | 3424.35 | $k9f2$ | 3.3289 |
| $k2d$ | 280.753 | $k9b2$ | 30.9967 |
| $k2f$ | 2.9131 | $km9$ | 5872.68 |
| $k2g$ | 0.267089 | $n9$ | 0.985466 |
| $k3a$ | 0.00137731 | $kbf$ | 0.01 |
| $k3b$ | 0.0987558 | $nC$ | 2.1e-06 |
| $k4a$ | 5790.17 | $INS_{offset}$ | 7 |
| $k4b$ | 34.7965 | $k_{gluin}$ | 2.14 |
| $k4c$ | 4.45581 | $p1$ | 0.179 |
| $k4e$ | 42.8395 | $p2$ | 4.48 |
| $k4f$ | 143.597 | $p3$ | 0.161 |
| $k4h$ | 0.536145 | $p4$ | 2.63 |
| $k5a1$ | 1.8423 | $k_{G6P}$ | 11495 |
| $k5a2$ | 0.055064 | $V_{G6Pmax}$ | 410 |
| $k5b$ | 24.826 | $k6f1$ | 2.65168 |
| $k5d$ | 1.06013 | $k5c$ | 0.0857515 |
| $km5$ | 2.64988 | | |

Table S1: Parameters of the cell level that where not changed. The values used where the same as in (Brännmark et al. 2013) for both the Topiramate Study and the Fast food study.

| Annotation | Value |
|---|---|
| $kIR$ | 0.0323845 |
| $kIRdeg$ | 1 |
| $kGLUT$ | 62.81266 |
| $kGLUTdeg$ | 1 |

Table S2: Parameters of the cell level that where changed. Fitted so that $IRtot$ and $GLUT4$ are at 100 % in steady state. The same values where used for both the Topiramate Study and the Fast food study.

| Annotation | Value used on Fast-food study data (estimated) | Value used on Topiramate study data (Herrgårdh et al. 2021) |
|---|---|---|
| $V_G$ | 1.98 | 1.88 |
| $k_1$ | 0.0613 | 0.065 |
| $k_2$ | 0.0779 | 0.079 |
| $G_b$ | 86.2 | 95 |
| $V_I$ | 0.0538 | 0.05 |
| $m_1$ | 0.208 | 0.19 |
| $m_2$ | 0.517 | 0.484 |
| $m_4$ | 0.192 | 0.194 |
| $m_5$ | 0.0274 | 0.0304 |
| $m_6$ | 0.5921 | 0.6471 |
| $HE_b$ | 0.660 | 0.6 |
| $I_b$ | 27.5 | 25 |
| $S_b$ | 1.83 | 1.8 |
| $k_{max}$ | 0.0525 | 0.0558 |
| $k_{min}$ | 0.00878 | 0.008 |
| $k_{abs}$ | 0.0624 | 0.057 |
| $k_{gri}$ | 0.0592 | 0.0558 |
| $f$ | 0.919 | 0.9 |
| $b$ | 0.741 | 0.82 |
| $dd$ | 0.0106 | 0.01 |
| $k_{p1}$ | 2.43 | 2.7 |
| $k_{p2}$ | 0.00209 | 0.0021 |
| $k_{p3}$ | 0.00968 | 0.009 |
| $k_{p4}$ | 0.0557 | 0.0618 |
| $k_i$ | 0.00714 | 0.0079 |
| $p_{2U}$ | 0.0316 | 0.0331 |
| $K$ | 2.49 | 2.3 |
| $alpha$ | 0.0480 | 0.05 |
| $beta$ | 0.101 | 0.11 |
| $gamma$ | 0.492 | 0.5 |
| $k_{e10}$ | 0.000458 | 0.0005 |
| $k_{e2}$ | 345 | 339 |
| $D$ | 67600 / 68110 | 101000 / 94759 |

Table S3: Parameters on the organ/tissue level that where set to different values for the different studies. For the Fast-food study, the parameters where estimated on initial values of glucose and insulin. For the Topiramate study, the values from (Herrgårdh et al. 2021) where used. The meal, $D$ (set to the same as the initial value of $Q_{sto}$), was set to 1 g/kg of body weight. The two values for D are for the meals simulated in the before/after scenarios respectively.

| Annotation | Value |
|---|---|
| $U_{ii}$ | 0.831 |
| $k2$ | 0.0429 |
| $k1$ | 0.0476 |
| $V_m$ | 0.881 |
| $V_{mx}$ | 0.0409 |
| $K_m$ | 476 |
| $V_l$ | 2.00 |
| $V_{lx}$ | 0.0439 |
| $K_l$ | 355 |
| $bf_b$ | 0 |
| $bradykinin$ | 1 |
| $be$ | 3 |
| $INS_b$ | 0.8549 |
| $p_{bf}$ | 5 |

Table S4: Parameters of the organ/tissue level that where kept at the same values for both the Fast food study and Topiramate study data. The values where come from (Herrgårdh et al. 2021).

| annotation | Value used on Topiramate study data (Herrgårdh et al. 2021) | Value used on Fast-food study data (estimated) |
|---|---|---|
| $dosage$ | 0 / 64 / 192 / 96 | 0 |
| $BWinit$ | 105.4 / 103.1 / 101 / 104.6 | 67.6 |
| $height$ | 166.5 / 166.2 / 166.1 / 166.8 | 175.6917 |
| $age$ | 43.6 / 45.1 / 44.7 / 45 | 27 |
| $RMRinit$ | 7189504 / 7054940 / 6971848 / 7133704 | 7102549 |
| $Ginit$ | 1.85 | 1.85 |
| $ECFinit$ | 17.34 / 16.96 / 16.6145 / 17.21 | 11.12 |
| $Finit$ | 52.12 / 50.44 / 48.59 / 51.49 | 12.7 |
| $Linit$ | 29.09 / 28.86 / 28.95 / 29.06 | 36.9348 |
| $ATinit$ | 0.1 | 0.1 |
| $EIrestriction$ | -600 | 3480 |
| $k$ | 0.010161196182943 | 0 |
| $lmax$ | 0.161411312363267 | 0 |
| $IC_{50}$ | 0.149118981539457 | 1 |
| $h2$ | 1 | 1 |
| $h1$ | 0.563148507725789 | 4.999973082721017 |
| $dEI_{ss}$ | 72.848728668834810 | 1.226464396357887e+03 |
| $t_{half}$ | 19.999999526293507 | 1.424047740519107 |

Table S5: Parameters of the whole-body level that where set to different values for the different studies. For the Fast-food study, the parameters where estimated on initial values of glucose and insulin. For both studies, the values where either taken from the data ($dosage$, $BWinit$, $height$, $age$, $Finit$, $Linit$, $EIrestriction$), if not present in data calculated based on the equations in (Hall et al. 2011) and data ($RMRinit$, $Ginit$, $ECFinit$, $Finit$, $Linit$, $ATinit$), or estimated to data from the Topiramate study ($k$, $lmax$, $IC_{50}$, and $h2$ to data for dosages 64 and 192, and $h1$, $dEI_{ss}$, and $t_{half}$ to placebo data). The different values for $dosage$, $BWinit$, $height$, $age$, $RMRinit$, $ECFinit$, $Finit$, $Linit$, and $EIrestriction$ are the data for the different dosages and their data sets. The meal, $D$ (set to the same as the initial value of $Q_{sto}$), was set to 1 g/kg of body weight. The two values are for the before/after scenarios respectively. See (Hall et al. 2011) for descriptions of parameters.

| Annotation | Value |
|---|---|
| $joules$ | 4183 |
| $fCIn$ | 0.4 |
| $PAE$ | 1.5 |
| $rhol$ | 7.6e6 |
| $rhof$ | 39.5e6 |
| $gf$ | 13000 |
| $gl$ | 92000 |
| $etal$ | 960000 |
| $etaf$ | 750000 |
| $rhog$ | 17.6e6 |
| $Na$ | 3.22 |
| $epNa$ | 3000 |
| $epCI$ | 4000 |
| $deltaNaDiet$ | 0 |
| $bTEF$ | 0.1 |
| $bAT$ | 0.14 |
| $tAT$ | 14 |
| $scale$ | 2.35 |
| $b_{CLGI}$ | 0.605258205407204 |
| $b_{EGP}$ | 1.142839561915420 |
| $b_{Ins}$ | 0.749598731223457 |
| $b_{IR}$ | 1.3696 |
| $b_{GLUT}$ | 1.5865 |
| $b_{diabetes}$ | 0.8962 |
| $delta$ | 30 |

Table S6: Parameters of the whole-body level that where kept at the same values for both the Fast food study and Topiramate study data. The values come from (Hall et al. 2011). See (Hall et al. 2011) for descriptions of parameters.

| Annotation | Value |
|---|---|
| $joules$ | 4183 |
| $CL$ | 1.21 |
| $V$ | 4.61 |
| $k10$ | 0.2625 |
| $Ka$ | 0.105 |
| $K23$ | 0.577 |
| $K32$ | 0.0586 |

Table S7: Parameters of the topiramate model used on the Topiramate study data. The values come from (Girgis et al. 2010)

| Annotation | Value used on Topiramate study data (Herrgårdh et al. 2021) | Value used on Fast-food study data (estimated) |
|---|---|---|
| $IRm(0)$ | 79.702633787375790 | 79.702633641196710 |
| $IRp(0)$ | 0.001616091202093 | 0.001616091199129 |
| $IRins(0)$ | 0.287638353419841 | 0.287638352892073 |
| $IRip(0)$ | 19.975821355446970 | 19.975821259258453 |
| $IRi(0)$ | 0.032384504361388 | 0.032384504291868 |
| $IRS1(0)$ | 0.636270043962264 | 0.636270048741165 |
| $IRS1p(0)$ | 0.004886920066483 | 0.004886920079528 |
| $IRS1p307(0)$ | 8.336225350721554 | 8.336225373343643 |
| $IRS1307(0)$ | 91.022683564690510 | 91.022683812180720 |
| $X(0)$ | 99.993271954939890 | 99.993271954921270 |
| $Xp(0)$ | 0.006815145060111 | 0.006815145078301 |
| $PKB(0)$ | 0.607375249849608 | 0.607375246877010 |
| $PKB308p(0)$ | 0.055433526447741 | 0.055433526415003 |
| $PKB473p(0)$ | 28.457706154130346 | 28.457706084935324 |
| $PKB308p473p(0)$ | 70.879375465504100 | 70.879375485218280 |
| $mTORC1(0)$ | 15.974509276998015 | 15.974509273264948 |
| $mTORC1a(0)$ | 84.025510723002180 | 84.025510726735000 |
| $mTORC2(0)$ | 38.229246047154106 | 38.229246160869980 |
| $mTORC2a(0)$ | 61.770764952846040 | 61.770764839130210 |
| $AS160(0)$ | 24.123630120570360 | 24.123630120162524 |
| $AS160p(0)$ | 75.876369879429820 | 75.876369879837410 |
| $GLUT4m(0)$ | 62.812699999999055 | 62.812699999999130 |
| $GLUT4(0)$ | 37.187401271425344 | 37.187401271225625 |
| $S6K(0)$ | 95.802072569195220 | 95.802072569021830 |
| $S6Kp(0)$ | 4.197900430804610 | 4.197900430977912 |
| $S6(0)$ | 68.925773402622870 | 68.925773401738680 |
| $S6p(0)$ | 31.074136597377260 | 31.074136598261480 |
| $Glu_{in}(0)$ | 6.219327923337478 | 5.491579474540069 |
| $G6P(0)$ | 0.005931193832426 | 0.005736832943166 |

Table S8: Initial values of cell level model. For the Topiramate study, these values where only used for the predictions (Fig. 5 E-G). These values where obtained through steady state simulation.

| Annotation | Value used on Topiramate study data (Herrgårdh et al. 2021) | Value used on Fast-food study data (estimated) |
|---|---|---|
| $G_p(0)$ | 179.7181945729903 | 166.8653475365416 |
| $G_t(0)$ | 134.5262549302064 | 120.2490760033025 |
| $I_l(0)$ | 5.683414265848025 | 5.327358533997125 |
| $I_p(0)$ | 1.592697213143252 | 1.566977929053884 |
| $Q_{sto1}(0)$ | 0 / 101000 | 0 |
| $Q_{sto2}(0)$ | 0 | 0 |
| $Q_{gut}(0)$ | 0 | 0 |
| $I_1(0)$ | 31.853944262865035 | 29.117931918456380 |
| $I_d(0)$ | 31.853944262865035 | 29.117931918456380 |
| $I_{po}(0)$ | 3.730852556413765 | 3.335612126317704 |
| $Y(0)$ | 0.065426278206883 | 0.187892188500337 |
| $INS_f(0)$ | 0 | 1.068266605322548e-19 |
| $INS(0)$ | 0 | 1.796308844088321 |

Table S9: Initial values of organ/tissue level model. For the Topiramate study, these values where only used for the predictions (Fig. 5 E-G). These values where obtained through steady state simulation, except for $Q_{sto1}(0)$ which was set to a meal corresponding to 1 g/kg body weight, or 0 if no meal.

| Annotation | Value used on Topiramate study data (Herrgårdh et al. 2021) | Value used on Fast-food study data (estimated) |
|---|---|---|
| $Gly(0)$ | 1.85 | 1.85 |
| $ECF(0)$ | 17.34 / 16.96 / 16.6145 / 17.21 | 11.1202 |
| $F(0)$ | 52.12 / 50.44 / 48.59 / 51.49 | 12.7 |
| $L(0)$ | 29.09 / 28.86 / 28.95 / 29.06 | 36.9348 |
| $AT(0)$ | 0.1 | 0.1 |

Table S10: Initial values of whole-body level model used in model training and validating. The different values for the Topiramate study correspond to the different dosages 0 / 64 / 192 / 96. For both studies, the values where either taken from the data ($F(0)$, and $L(0)$), or if not present in data calculated based on the equations in (Hall et al. 2011) and data ($RMR(0)$, $Gly(0)$, $ECF(0)$, $F(0)$, $L(0)$, $AT(0)$).

| Annotation | Values used for dosage 64 mg/day | Values used for dosage (192 mg/day) | Values used for dosage 96 mg/day (validation) |
|---|---|---|---|
| $dosage(0)$ | 16 / 32 / 64 | 16 / 32 / 64 / 96 / 128 / 192 | 16 / 32 / 64 / 96 |
| $C(0)$ | 0 | 0 | 0 |
| $y3(0)$ | 0 | 0 | 0 |

Table S11: Initial values of topiramate model used in model training and validating for Topiramate study. The dosage of topiramate is given as an initial value of dosage, and the different values for each dosage represents the escalation of dosage taken seen in the data in (Bray et al. 2003).

| **Full-length name** | **Abbrevation** |
|---|---|
| Type 2 diabetes | T2D |
| Insulin receptor | IR |
| Insulin receptor substrate 1 | IRS1 |
| Protein kinase B | PKB |
| Endogenous glucose production | EGP |
| Insulin in the portal vein | Ipo |
| Ordinary differential equations | ODEs |
| Food and drug administration | FDA |
| Fat mass | FM |
| Fat free mass | FFM |
| Energy intake | EI |
| Energy expenditure | EE |
| Body weight | BW |
| Phosphorylated PKB | PKB308-p |
| Phosphorylated IRS1 | IRS1-p |

Table S12: Abbreviations.
